# A bistable UV-sensitive opsin from a reef building coral showing a switchable and tunable regulation of Gs-signaling by different wavelengths of light

**DOI:** 10.64898/2026.07.01.735937

**Authors:** Yusuke Sakai, Akinari Sakayori, Tomoki Kawaguchi, Kento Takano, Keita Sato, Keiichi Kojima, Hideyo Ohuchi, Hisao Tsukamoto

## Abstract

Cnidarians possess large number of opsins in their genomes for their various photoreceptive functions. In particular, they uniquely possess Gs-coupled opsins that induce intracellular cAMP accumulation in a light-dependent manner. These Gs-coupled opsins, cnidopsins, are powerful optogenetic tools manipulating cAMP-dependent cellular responses. In this study, we characterized a cnidopsin, named as AtCnidop3a, from the coral *Acropora tenuis* as a Gs-coupled and UV-sensitive bistable pigment. This cnidopsin showed a large spectral shift upon activation from absorption maxima from 395 nm to 560 nm, and the resting and activated states are interconvertible by illumination with UV (or violet) and orange light. The activated state efficiently activated Gs proteins and elevated intracellular cAMP levels in mammalian cultured cells. To engineer the opsin mutant that can be turned on and off upon long wavelength light illumination by utilizing the large spectral separation, negatively charged amino acids were introduced near the retinal Schiff base region. Among tested opsin mutants, the Y113^3.28^E mutant is capable of being activated by green light unlike the wild-type while retaining the property of being inactivated by orange light like the wild-type, indicating successful conversion of the opsin to a visible light sensitive bistable pigment. The visible light-induced cAMP regulation of the Y113^3.28^E mutant was enhanced by an additional L94^2.61^G substitution. Our characterization and engineering of the cnidopsin revealed functional diversity of cnidarian opsins and its potential utility as optogenetic tools regulating Gs-dependent physiological responses.

## Introduction

Reef-building corals use environmental light stimuli to regulate their physiologies, including larval behaviour^1–3^ and spawning^4–6^. For example, larvae of a Caribbean coral *Porites astreoides* swim to avoid UV irradiation^7^, and larvae of a reef-building coral *Acropora tenuis* stop swimming upon attenuation of short-wavelength light^3^. These behaviours may be important to settle to the sea floor with less light damage. Such physiological response to light starts with photon catches by photoreceptive proteins including opsins, which are primary photoreceptive proteins in animals.

Opsins are light-sensitive G protein-coupled receptors (GPCRs) having a seven-transmembrane α-helical structure and stimulate heterotrimeric G protein complexes, followed by the activation of downstream intracellular signaling cascades^8,9^. Animals belonging to the phylum Cnidaria including corals are known to possess multiple opsin genes. Cnidarians uniquely possess Gs-coupled opsins and these opsins referred to as cnidops^10^ or cnidopsins^11^. Cnidopsins were first identified from Hydra^10^, and they were later found in a broad range of cnidarian animals such as jellyfishes^12–14^, sea anemones^15^, and corals^16–18^, too. Previous studies on various cnidopsins revealed their functional diversity.

Among various cnidopsins, the box jellyfish cnidopsin (JellyOP) showed a green-light absorption [absorption maximum (λmax) at 500 nm] in the spectroscopic analysis^12^. A subsequent study has shown that cnidopsins (acropsins 1, 2, and 6) from a reef-building coral, *Acropora millepora*, have an ability to absorb visible light and cause light-dependent increase in cAMP levels in mammalian cultured cells^17^. The study estimated the spectral sensitivities of the two cnidopsins with their maximum sensitivities at 472 nm (acropsin 1) and 476 nm (acropsin 6) using heterologous action spectroscopy, which is a method to estimate spectral sensitivity based on light wavelength- and intensity-dependent cAMP increases of opsin-expressing cultured cells. The spectral sensitivity of acropsin 2 was also estimated in a similar way to be the λmax at 471 nm. Another spectroscopic study characterized cnidopsins from a different reef-building coral species *Acropora tenuis* and showed the AtCnidop6 and AtCnidop7 are blue- and UV- sensitive bistable pigments, respectively^18^.

The Gs-coupling and green-light sensitivity of JellyOP has been utilized to optogenetically regulate Gs-dependent cellular responses^19,20^, in particular intracellular cAMP elevation via activation of adenylyl cyclase by Gs. The spectral diversity in cnidopsins indicates their potential utility as optogenetic tools modulating Gs-dependent physiological responses by various color of light stimulations.

In this study, we show absorption spectrum of purified pigments of a cnidopsin from the reef-building coral *Acropora tenuis* (hereafter, the cnidopsins is referred as AtCnidop3a) (see Fig. 1A), showing that the opsin has a sensitivity to ultraviolet (UV) light. Notably, based on the spectroscopic measurement, we demonstrate that AtCnidop3a exhibits a bistable nature — interconvertible photoreaction between the dark state and the active state by irradiation of different wavelengths of light. In consistent with the spectroscopic feature, AtCnidop3a shows increase and decrease of Gs activity and intracellular cAMP levels repeatedly by repetitive UV-and orange-light irradiations. Furthermore, we introduced amino acid substitutions into AtCnidop3a, and converted the opsin to green-light activated and orange light deactivated bistable opsin. Based on the insights and functional modification of characteristics in AtCnidop3a, we discuss its potential as a new Gs-coupled optogenetic tool for switchable regulation of intracellular cAMP levels by different wavelengths of light.

**Fig. 1.**
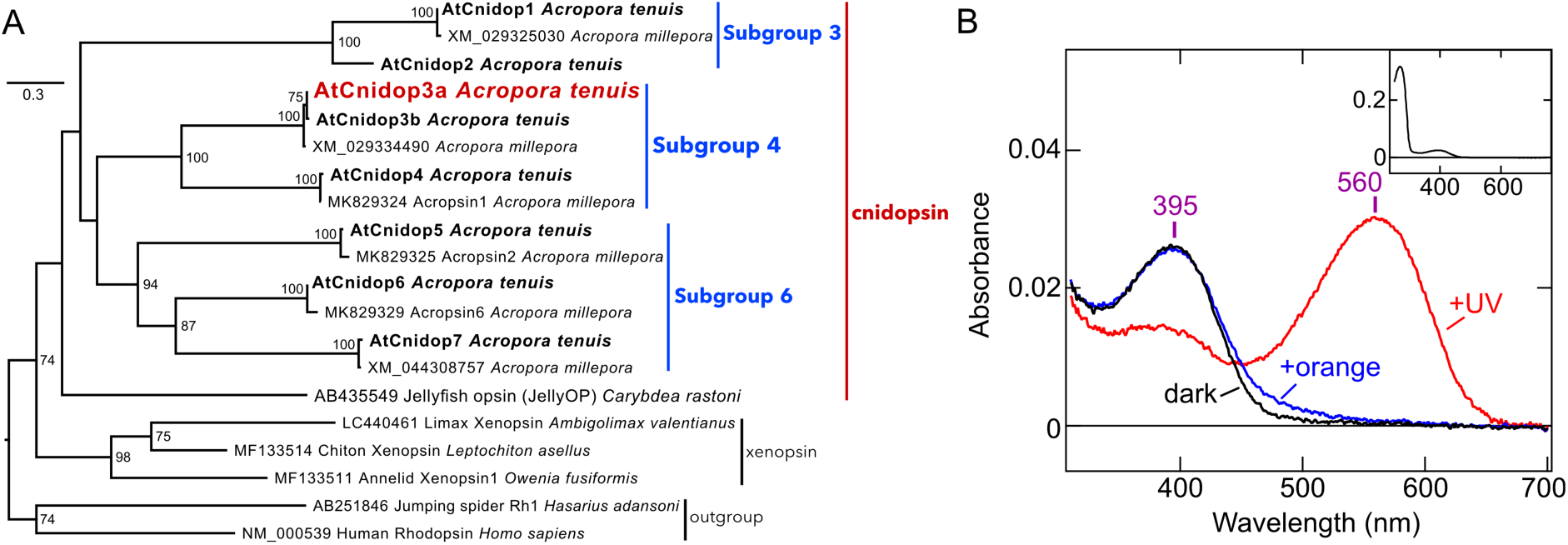
Phylogenetic relationship and spectral properties of AtCnidop3a. A, Molecular phylogenetic tree of cnidopsins. The maximum likelihood tree was reconstructed based on amino acid sequences of several coral cnidopsins, JellyOP, human rhodopsin, jumping spider Rh1, and xenopsins (which have been reported as the opsin group most closely related to cnidopsins^15,57^). *Acropora tenuis* Cnidopsins (AtCnidops, GenBank accession numbers: LC844924 – LC844931) are shown as bold letters and AtCnidop3a is highlighted by red. Cnidopsin subgroups based on the previous study^18^ are indicated as blue letters. Numbers at the nodes show support values examined by 1000 bootstrap samplings (≥ 70% were indicated). Scale bar shows 0.3 substitutions per site. B, Absorption spectra of purified AtCnidop3a WT. Spectra in the dark (*black*), after UV illumination (*red*), and after subsequent orange light illumination (*blue*) are shown. In the dark, λ_max_ value is 395 nm, and after UV light illumination, the value is shifted to 560 nm. (*Inset*) Absorption spectrum of AtCnidop3a in the dark in a wider wavelength range.

## Materials and Methods

### Phylogenetic tree inference

Amino acid sequences of opsins were aligned using MAFFT^21^ and the multiple alignment was trimmed using TrimAl^22^ with the “gappyout” function. The maximum likelihood (ML) tree was generated assuming the LG+I+G4+FC model of protein evolution and support for the nodes was examined by 1000 bootstrap iterations. The model selection and ML tree reconstruction was performed using ModelTest-NG v0.2.0^23^ and RAxML-NG v1.2.0^24^, respectively.

### Constructs

The cDNAs of JellyOP WT^12^ and AtCnidop3a WT^18^ and mutants were tagged with the 1D4 sequence (ETSQVAPA) on their C-termini and inserted into the *Eco*RI/*Not*I site in an expression vector pMT using in-Fusion HD (TAKARA, Japan). The RIC8B isoform2/pCAGGS was generously gifted by Dr. Asuka Inoue (Kyoto University). The cDNAs of Nluc inserted human GsαL and Venus fused human Gγ1 were constructed and inserted into the pMT vector according previous BRET studies^25,26^. The cDNA of human Gβ1 in the pcDNA3.1 was obtained from addgene (#140987)^25^. The amino acid sequences of the Nluc inserted human GsαL and the Venus fused human Gγ1 are shown in Supplemental Data.

### Protein expression and purification of AtCnidop3a

AtCnidop3a WT and L94^2.61^G/Y113^3.28^E mutant were transiently expressed in COS-1 cells (10 plates), and the cells were harvested 48 h after transfection as described previously^27,28^. The collected cells were incubated with 11-*cis*-retinal overnight, and membrane proteins were solubilized with 1.25 % DDM (Dojindo, Japan), 20 mM HEPES, 140 mM NaCl, 0.25 % cholesterol hemisuccinate (Sigma-Aldrich, St. Louis, MO) 25 mM Tris, 10 % glycerol, pH 7.0. The solubilized materials were mixed with 1D4-agarose overnight, and the mixture was transferred into Bio-Spin columns (Bio-rad, Hercules, CA). The columns were washed with 0.05 % DDM, 2 mM ATP, 1 M NaCl, 3 mM MgCl_2_, 0.01 % cholesterol hemisuccinate, 1 mM Tris, 10 % glycerol in PBS, and subsequently washed with 0.05 % DDM, 140 mM NaCl, 0.01 % cholesterol hemisuccinate, 1 mM Tris, 10 % glycerol, 20 mM HEPES, pH 7 (buffer A). The 1D4 tagged pigments were eluted with buffer A containing 0.45 mg/mL 1D4 peptide (TETSQVAPA) (TOYOBO, Japan).

### UV-Vis Spectroscopy

Absorption spectra of purified opsins were recorded with a Shimadzu UV-2600 spectrophotometer (Shimadzu, Japan). The samples were kept at 10 °C. An optical interference filter YIF-BP340-390S (Sigma-Koki, Japan), which transmits light around 370 nm, was used for illumination of AtCnidop3a WT with UV light (duration: 1 min, light intensity after the filter: ∼9 mW/cm^2^, light source: 250 W halogen lamp). In order to convert the photoproduct to the resting state, a longpass filter SCF-50S-58O (Sigma-Koki), which transmits light longer than 580 nm, was used (duration: 30 sec, light intensity after the filter: ∼140 mW/cm^2^, light source: 250 W halogen lamp) as orange light.

### Transfection of constructs for GloSensor and BRET-based Gs Protein Dissociation Assays

Opsins, GloSensor assay sensor (coded by pGlo-22F), and BRET sensor G protein subunits were transiently expressed in COS-1 cells using polyethyleneimine as described previously^27,29^. For the BRET-based Gs dissociation assay, each well of 96-well assay plate (Corning, Kennebunk, ME) was transfected with 50 ng opsin plasmid, 25 ng NLuc inserted GsαL plasmid, 50 ng Gβ_1_ plasmid, 50 ng Venus fused Gγ_1_ plasmid, 50 ng RIC8B plasmid, and 500 ng polyethyleneimine in 25 μL Opti-mem (Gibco) and 75 μL D-MEM (Wako) containing 10 % (vol/vol) FBS, 100 units/mL penicillin, and 100 μg/mL streptomycin. For the GloSensor assay, each well of 96-well assay plate (Corning) was transfected with 50 ng opsin plasmid, 50 ng pGlo-22F plasmid (Promega, Madison, WI), and 500 ng polyethyleneimine in 25 μL Opti-mem (Gibco, Waltham, MA) and 75 μL D-MEM (Wako, Japan) containing 10 % (v/v) FBS, 100 units/mL penicillin, and 100 μg/mL streptomycin.

### GloSensor Assay

The transfected COS-1 cells were incubated at 37 °C, 5 % CO_2_ for 2 days, and the medium was aspirated, followed by addition of HBSS (145 mM NaCl, 10 mM D-glucose, 5 mM KCl, 1 mM MgCl_2_, 1.7 mM CaCl_2_, 1.5 mM NaHCO_3_, 10 mM HEPES, pH 7.4) containing 1 μM 11-*cis*-retinal and 2 % (vol/vol) GloSensor cAMP reagent stock solution. Then, the cells were incubated at room temperature for ∼2 h to stabilize luminescence levels. Luminescence was measured using GM-2000 or GM-3510 microplate reader (Promega). Luminescence level of each well was measured every 30 sec with integration time of 0.9 sec. To illuminate opsin, luminescence measurement was interrupted, the plate was ejected, and the plate was illuminated by blue or orange light. UV, green, and orange light sources are 390 nm (duration: 1 min, light intensity, ∼0.1 mW/cm^2^ or duration: 5 sec, light intensity, ∼10 mW/cm^2^), 520 nm (duration: 1 min, light intensity, ∼0.3 mW/cm^2^) and 600 nm (duration: 1 min, light intensity, ∼0.4 mW/cm^2^ or duration: 5 sec, light intensity, ∼30 mW/cm^2^) from Opto-spectrum generator L12194 (Hamamatsu, Japan), respectively. After illumination, luminescence measurement was resumed. The measured luminescence levels were normalized to the level at the starting point (time = 0 min).

### Establishment of HEK293T Gs/Gi/Gq KO cells

The KO cells were generated on the basis of *GNAQ*/*GNA11*/*GNA14*/*GNA15* quadruple-knockout HEK293T cells established in the previous study^30^. Target sites for genome editing were chosen using the web tool CRISPRdirect^31^ (https://crispr.dbcls.jp/) to minimize the possibility of off-target cleavage. Information on the chosen target sequences is summarized in Supplemental Fig. S3 and Supplemental Table S1. HEK293T cells were seeded into 24-well plates in Dulbecco’s modified Eagle’s medium/F-12 (FUJIFILM Wako, Japan) supplemented with 10% FBS. One day later, the cells were transfected with a SpCas9-2A-Puro expression plasmid and an sgRNA expression plasmid using HilyMax (DOJINDO, Japan) according to the manufacturer’s instructions. Six to eight hours after transfection, the medium was replaced with fresh medium. Twenty-four hours after transfection, puromycin was added at 5 μg/ml. Twenty-four hours after puromycin addition, live cells were trypsinized, counted, and reseeded onto 10-cm culture dishes at 100 to 300 cells per dish. After about 15 days, clonal colonies on the culture dishes were picked by scratching and transferred to 24-well plates. To determine whether the target Gα gene had been knocked out, each clone was screened by PCR of genomic DNA. In principle, two genes were targeted at a time, and transfection, cloning, and screening were repeated sequentially. Because PCR screening sometimes suggested that wild-type sequences remained, some genes were retargeted by repeated transfection of the targeting sgRNA plasmid. The genes were targeted in the following order: *GNAO*/*GNAZ*, *GNAI1*/*GNAI2*, *GNAI3*/*GNAT1*, *GNAT2*/*GNAT3*, *GNAT2*/*GNAS*, *GNAS*/*GNAL*, and finally *GNAT2* again. Supplemental Fig. S3 shows the genotyping results for the final clones, which were determined by PCR amplification of the genomic regions of the individual genes, sequencing of the PCR products, and outsourced whole-genome sequencing analysis (Macrogen, Japan). The primer sequences used for genotyping are listed in Supplemental Table S2. Whole-genome sequencing raw reads for the final KO clone were deposited in the DDBJ Sequence Read Archive under BioProject accession [PRJDB42188].

### BRET-based Gs Protein Dissociation Assay

The BRET-based Gs dissociation assay in this study was conducted according to previous BRET studies^25,26^. The transfected COS-1 cells were incubated at 37 °C, 5 % CO_2_ for 1 day, and the medium was aspirated, followed by addition of HBSS containing 1 μM 11-*cis*-retinal and 0.2 % (vol/vol) Nano-Glo Luciferase Assay reagent (Promega). Then, the cells were incubated at room temperature for 15 min to stabilize luminescence levels. Doner luminescence and acceptor fluorescence were measured using GM-3510 microplate reader equipped with BRET/FRET Upgrade GM-3570 (Promega). Luminescence level of each well was measured every 30 sec with integration time of 0.5 sec. Nluc (doner) luminescence was selected using an interference filter transmitting 450 nm (8 nm bandpass) (Promega), and Venus (acceptor) fluorescence was selected using a interference filter transmitting longer than 525 nm (Edmund Optics, Barrington, NJ). The BRET ratio (A/D ratio) was calculated as the acceptor Nluc fluorescence divided by the doner Venus luminescence. The ΔBRET values were calculated as the BRET ratios at respective data points minus the BRET value at the starting point (time = 0 min). To illuminate opsin, luminescence measurement was interrupted, the plate was ejected, and the plate was illuminated by blue or orange light. UV, green, and orange light sources are 420 nm (duration: 30 sec, light intensity, ∼0.4 mW/cm^2^), 520 nm (duration: 30 sec, light intensity, ∼0.3 mW/cm^2^) and 600 nm (duration: 30 sec, light intensity, ∼0.4 mW/cm^2^) from Opto-spectrum generator L12194 (Hamamatsu, Japan), respectively. The measured BRET ratios were normalized to the level at the starting point (time = 0 min).

## Results

### Phylogenetic relationship and spectral properties of AtCnidop3a

Cnidopsins from the coral *Acropora tenuis* (AtCnidops) are classified into several cnidopsin subgroups. The cnidopsin subgroup 3, 4, and 6 are shown in Fig. 1A. A previous study characterized AtCnidop6 and AtCnidiop7 belonging to the same cnidopsin subgroup (Subgroup 6 in Fig. 1A) as blue and UV light-sensitive bistable opsin, respectively^18^. AtCnidop3a belongs to a different cnidopsin subgroup (Subgroup 4 in Fig. 1A) from AtCnidop6 and AtCnidiop7. In the previous study, the authors tried but failed to characterize AtCnidop4 in the subgroups 4, due to poor expression in cultured cells^18^. In the present study, we tried to express a 1D4-tagged AtCnidop3a in mammalian cultured cells (COS-1 cells) and successfully purified the photopigment after reconstitution with retinal, extraction, and immuno-column chromatography (see “Materials and Methods”).

The purified AtCnidop3a absorbed UV light in the dark and its λmax was at 395 nm (Fig. 1B, black curve). To investigate the photoreactions of the UV-sensitive AtCnidop3a, we illuminated the opsin with UV light and measured the absorption spectra after the light irradiation. AtCnidop3a showed a large red-shift of the absorbance to the visible region (λmax of 560 nm) upon the first irradiation with UV light (Fig. 1B, red curve), and the absorption spectra reverted to the original dark state having a single peak at ∼395 nm by second irradiation with orange light (>580 nm) to the photoproducts (Fig. 1B, blue curve). Repeated irradiations to AtCnidop3a with same wavelengths of light (UV and >580 nm) recreated the results such that the increase in the absorbance around 560 nm upon UV-light irradiation and the absorbance reverted to the similar curve upon orange-light irradiation (Supplemental Fig. S1). The repeated and interconvertible photoreaction indicate that AtCnidop3a is a typical bistable pigment. On the other hand, the absorption spectra after orange light irradiation to the photoproducts did not completely match to the original dark spectra (Supplemental Fig. S1) — having a broadened bandwidth which might be derived from degradation by the light irradiations.

Within our knowledge, the >160-nm red-shift in AtCnidop3a upon light illumination is the largest red-shift in bistable opsins. For example, UV light-sensitive bistable ciliary (Gi/o-coupled) opsins lamprey parapinopsin and *Platynereis* c-opsin1 showed ∼130-nm and ∼110-nm red-shifts, respectively^32,33^. In addition, a recent study characterizing rhabdomeric (Gq-coupled) opsins from mantis shrimp reported that some blue-sensitive opsins (NOM5 and NOM6) showed ∼100-nm red-shifts upon blue-light illumination^34^. The >160-nm red-shift of AtCnidop3a exceeds these reported values in other bistable opsins.

### Gs-coupling and cAMP response of AtCnidop3a

The bistable photoreaction of AtCnidop3a suggested that the opsin can be activated by UV light and deactivated by orange light. Since cnidopsins have been reported to be coupled with Gs-type G proteins^12,17,19,35^, we next assessed light dependent-Gs activation and deactivation by AtCnidop3a. We employed a BRET-based assay to investigate coupling between AtCnidop3a and Gs^25,26^. In this assay, the BRET donor luciferase Nluc and the BRET acceptor fluorescent protein Venus were fused to Gs alpha subunit (Gsα) and gamma subunit (Gγ), respectively (see “Materials and Methods”) (Fig. 2A). The Gs activation (dissociation of Gsα and Gβγ) was detected as decrease of BRET efficiency (negative ΔBRET value) and deactivation (association of Gsα and Gβγ) was detected as recovery of the ΔBRET signal. In COS-1 cells, no obvious changes in BRET signals were observed upon illumination of AtCnidop3a (Supplemental Fig. S2). We thought endogenous Gsα proteins without Nluc insertion interfered the BRET signals, and did the BRET assay using HEK293T cells lacking Gsα subunits. Then we used HEK293T Gs/Gq/Gi KO cells, in which Gsα, Golfα, Giα1, Giα2, Giα3, Goα, Gzα, Gtα1, Gtα2, Ggustα, Gqα, G11α, G14α, and G15α genes are knocked out (Supplemental Fig. S3 and Supplemental Table S1, also see “Materials and Methods”). In the KO cells, we detected sustained decrease of BRET signals after stimulation with violet light (420 nm, “+V”) and recovery of the signals upon subsequent orange light (600 nm, “+O”) illumination (Fig. 2B). To the best of our knowledge, this result is the first evidence to directly assess activation and deactivation of Gs caused by a cnidopsin. In contrast, visible light-sensitive JellyOP activates Gs either upon violet- or orange-light illumination, and the activity spontaneously decayed as reported in previous studies^19,36^ (Fig. 2C). These results clearly proved that AtCnidop3a is a bistable Gs-coupled opsin. In AtCnidop3a, the 395-nm species (black line in Fig. 1B) is the resting state and the 560-nm species (red line in Fig. 1B) is the stable activated state that is coupled with Gs proteins.

**Fig. 2.**
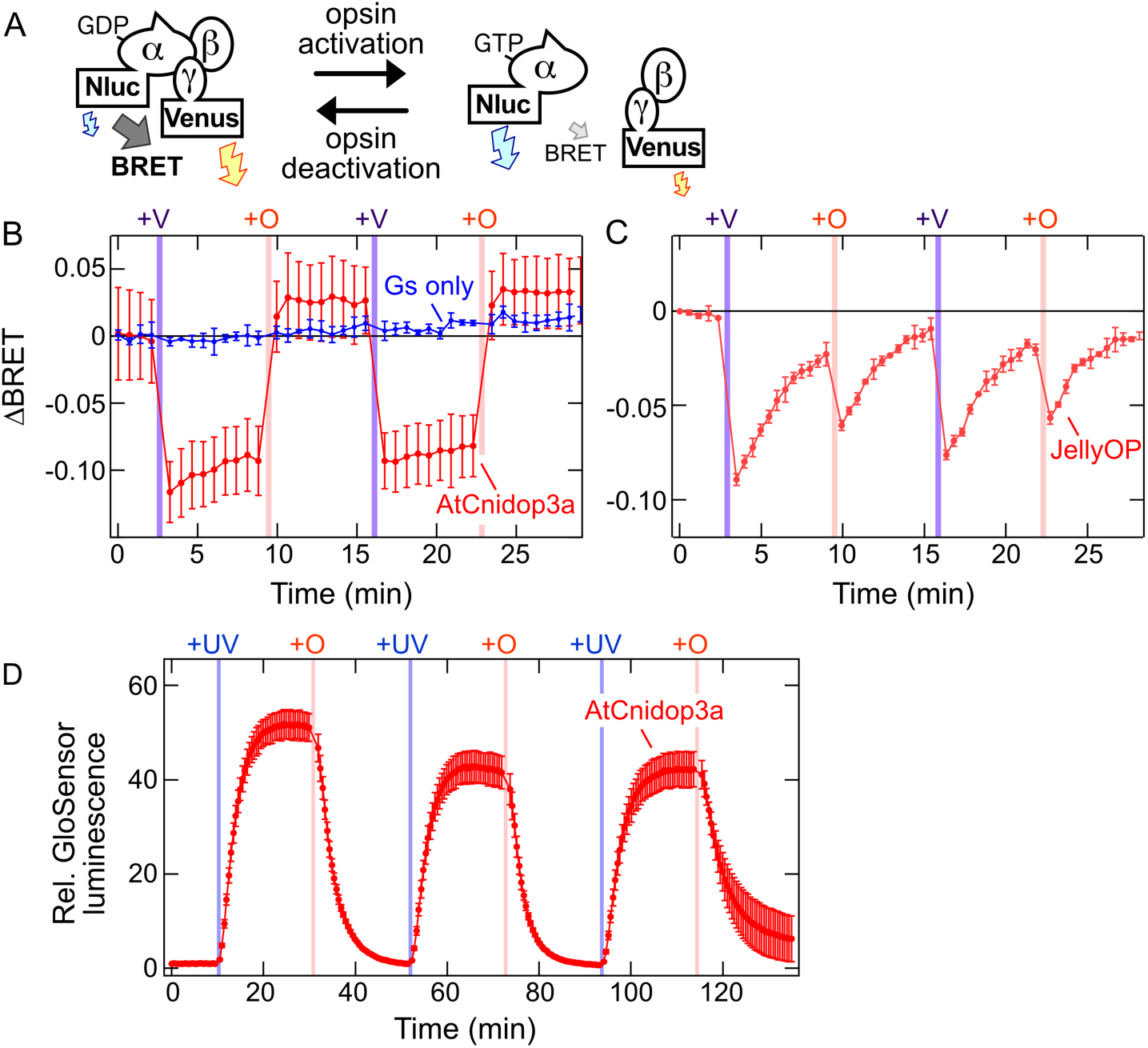
Activation and deactivation of AtCnidop3a. A, Schematic representation of the BRET-based Gs dissociation assay. In this BRET assay, BRET donor Nluc is fused into Gsα and BRET acceptor Venus is fused into Gγ. Activated opsin induces dissociation of Gα and Gβγ, leading to increases in the Venus fluorescence and the decrease in the BRET efficiency. When the opsin is deactivated, Gα and Gβγ (re)associate into the Gαβγ trimer leading to decrease in the Venus fluorescence and increase in the BRET efficiency. The detail is described in “Materials and Methods”. B, Light-induced changes in the BRET efficiencies (ΔBRET values) luminescence of HEK293T Gs/Gq/Gi KO cells expressing AtCnidop3a WT (*red*) and without the opsin (*blue*). C, Light-induced changes in the ΔBRET values of HEK293T Gs/Gq/Gi KO cells expressing JellyOP (*red*). ΔBRET time courses upon violet- (“+V”) and orange- (“+O”) light illuminations are shown. D, Light-induced changes in cAMP biosensor (GloSensor) luminescence of COS-1 cells expressing AtCnidop3a WT. Luminescence levels are normalized to the value at the starting point (time = 0 min). UV (“+UV”) and orange (“+O”) bars indicate UV and orange illuminations, respectively. In panels B, C, and D, Error bars indicate the SD values (n= 3).

Some Gs-coupled cnidopsins can cause increase in intracellular cAMP upon light illumination^17,20,35,36^. To evaluate the downstream signalling of AtCnidop3a, we measured light-dependent change in the cAMP level in the opsin-expressing COS-1 cells by the GloSensor cAMP assay^37^. We observed clear and sustained cAMP increase after 390 nm-light irradiation (“+UV”) (Fig. 2D), indicating AtCnnidop3a had a basic property as Gs-coupled opsin like other jellyfish^12,35,36^ and coral^17^ cnidopsins. Moreover, we found that the UV-induced cAMP increases were shut off by subsequent irradiation with orange (600 nm, “+O”) light, and the light-induced cAMP increase and decrease can be repeated (Fig. 2D), whereas cAMP responses by JellyOP is transient and spontaneously shut off^36^. These results indicated that AtCnidop3a can modulate intracellular cAMP levels bidirectionally via UV (or violet) light-dependent activation and orange light-dependent deactivation of Gs.

### Functional conversion of AtCnidop3a to a visible light-sensitive bistable pigment by Y113^3.28^E substitution

In the resting state, AtCnidop3a possesses λmax in UV region (395 nm) (Fig. 1B), suggesting that the retinal Schiff base linkage is deprotonated. In agreement with this speculation, the amino acid residue at position 94^2.61^ [according to bovine rhodopsin numbering system with Ballesteros/Weinstein (GPCRdb) numbering as superscript, see Supplemental Fig. S4] is occupied with leucine (Leu-94^2.61^) without negative charge on the side chain (see Fig. 3A), whereas green-light sensitive JellyOP possesses Glu-94^2.61^ as the counterion to stabilize the protonated retinal Schiff base^35^. Therefore, we speculated that AtCnidop3a lacks the counterion for the retinal Schiff base in the UV-sensitive resting state. On the other hand, the Schiff base of the activated state should be protonated, allowing the opsin to show the largely red-shifted λmax at 560 nm (Fig. 1B). Previous studies reported that spectral properties of opsins are somewhat independently tuned between the resting and activated states^27,38–40^, suggesting that introduction of artificial counterion can convert the resting state to visible light-sensitive while the activated state is intact. In other words, introducing the counterion into AtCnidop3a could make the opsin to be activated and deactivated by long wavelength light illuminations. In this context, we tried to engineer the opsin mutant that is activated by green light (520 nm) unlike the wild-type (WT) and deactivated by orange light (600 nm) like WT.

**Fig. 3.**
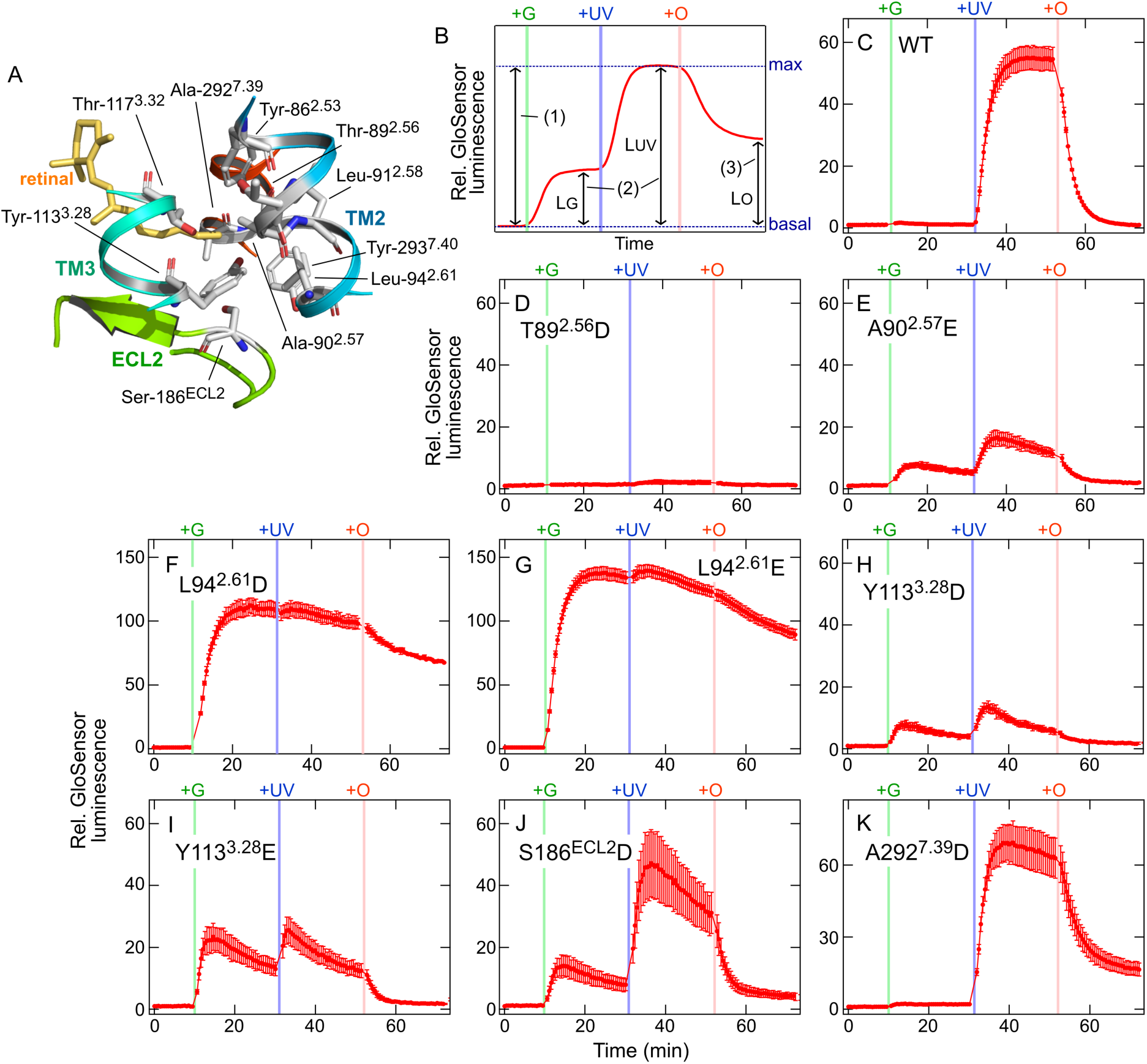
Functional conversion of AtCnidop3a from UV-sensitive to visible light-sensitive bistable opsin by site-directed mutagenesis. A, Arrangement of the amino acid residues at which we introduced amino acid substitution in this study in the AlphaFold2-predicted AtCnidop3a structure^41^. The 11-*cis*-retinal molecule is adopted from the crystal structure of bovine rhodopsin (PDB ID: 1U19)^58^. The structural models are prepared using PyMOL (https://pymol.org/). B, Schematic representation of the parameters of GloSensor signals in selection of visible light-sensitive bistable AtCnidop3a mutants. “L_G_”: the luminescence changes upon the first green light illumination. “L_UV_”: the luminescence changes upon the second UV light illumination. “L_O_”: the luminescence levels after the third orange-light illumination. “(1)”, “(2)”, and “(3)” indicate three criteria to evaluate photoresponses of the tested mutants (see the main text). C – K, Light-induced changes in cAMP biosensor (GloSensor) luminescence of COS-1 cells expressing AtCnidop3a WT (C), T89^2.56^D (D), A90^2.57^E (E), L94^2.61^D (F), L94^2.61^E (G), Y113^3.28^D (H), Y113^3.28^E (I), S186^ECL2^D (J), and A292^7.39^D (K). Luminescence levels are normalized to the value at the starting point (time = 0 min). Green (“G”), UV (“+UV”) and orange (“+O”) bars indicate Green, UV, and orange illuminations, respectively. Error bars indicate the SD values (n= 3). Summary of photoresponses of all tested AtCnidopsin3a single mutants are indicated in Table 1.

**Table 1.** Light-dependent GloSensor responses of AtCnidop3a mutants.

| mutant | L<sub>G</sub> | L<sub>UV</sub> | L<sub>O</sub> |
| --- | --- | --- | --- |
| WT | 1.72±0.03 | 55.1±4.00 | 0.97±0.04 |
| Y86<sup>2.53</sup>D | 2.31±0.54 | 3.19±0.20 | 1.19±0.10 |
| Y86<sup>2.53</sup>E | 2.19±0.08 | 7.14±0.50 | 2.32±0.07 |
| T89<sup>2.56</sup>D | 1.57±0.09 | 2.40±0.62 | 1.19±0.10 |
| T89<sup>2.56</sup>E | 2.11±0.20 | 2.86±0.26 | 1.54±0.15 |
| A90<sup>2.57</sup>D | 3.16±0.28 | 4.16±0.07 | 3.27±0.24 |
| A90<sup>2.57</sup>E | 8.01±0.55 | 16.7±2.15 | 1.93±0.13 |
| L91<sup>2.58</sup>D | 1.36±0.16 | 5.06±0.25 | 1.35±0.27 |
| L91<sup>2.58</sup>E | 1.68±0.21 | 13.1±3.07 | 1.71±0.32 |
| L94<sup>2.61</sup>D | 112±5.82 | 110±6.18 | 67.7±0.26 |
| L94<sup>2.61</sup>E | 138±4.80 | 140±4.94 | 89.6±4.12 |
| Y113<sup>3.28</sup>D | 8.01±1.30 | 13.7±1.82 | 1.66±0.18 |
| Y113<sup>3.28</sup>E | 23.4±4.36 | 25.6±4.22 | 1.73±0.16 |
| T117<sup>3.32</sup>D | 4.94±1.08 | 18.1±4.80 | 9.00±2.65 |
| T117<sup>3.28</sup>E | 127±10.2 | 163±14.3 | 149±12.1 |
| S186<sup>ECL2</sup>D | 14.0±3.23 | 47.2±10.8 | 4.02±1.14 |
| S186<sup>ECL2</sup>E | 1.61±0.22 | 35.3±6.27 | 7.07±1.78 |
| A292<sup>7.39</sup>D | 2.14±0.02 | 69.4±9.07 | 16.6±2.73 |
| A292<sup>7.39</sup>E | 4.30±0.18 | 92.5±7.17 | 23.1±2.71 |
| Y293<sup>7.40</sup>D | 1.41±0.16 | 3.25±0.37 | 1.28±0.18 |
| Y293<sup>7.40</sup>E | 1.35±0.07 | 7.39±0.44 | 1.53±0.05 |

Based on the AlphaFold2^41^ predicted structure of AtCnidop3a (Fig. 3A), we picked up 10 positions 86^2.53^, 89^2.56^, 90^2.57^, 91^2.58^, 94^2.61^, 113^3.28^, 117^3.32^, 186^ECL2^, 292^7.39^, and 293^7.40^ (Supplemental Fig. S4), all of which would be located near the retinal Schiff base, to introduce an artificial counterion. Previous studies on other opsins have reported that native or artificially-introduced Glu or Asp residues at position 90^2.57^, 94^2.61^, 113^3.28^, 117^3.32^, 186^ECL2^, or 292^7.39^ can function as counterion in their resting states^35,42–47^. In AtCnidop3a, we introduced Glu or Asp residue at each position and tested visible-light sensitivity of total 20 mutants using GloSensor assay (see Table 1).

In order to select the opsin mutants that is activated by green light and deactivated by orange light, we illuminated the opsin-expressing cells with three distinct wavelengths of light in the following order: 1st green light (520 nm, “+G”), 2nd UV light (390 nm, “+UV”), and 3rd orange light (600 nm, “+O”) with 20 min interval between light illuminations (see Fig. 3B). We monitored cAMP changes based on GloSensor luminescence signals, and evaluated their photoresponses on three criteria as follows: (1) maximal luminescence changes are greater than 20-fold of the basal luminescence to ensure sufficient signal transduction (light-dependent cAMP elevation), (2) the luminescence changes upon the first green light illumination (L_G_) are comparable to those after the second UV-light illumination (L_UV_), showing that the second UV illumination does not cause further activation of the opsin mutant, to ensure efficient green light sensitivity, (3) the luminescence levels after the third orange-light illumination (L_O_) decay to less than 3-fold of the basal luminescence to ensure sufficient light-dependent deactivation (Fig. 3B). For example, UV-sensitive AtCnidop3a WT met the criteria 1 (maximal response was ∼55-fold) and 3 (L_O_ was ∼1-fold) but not the criterion 2 (L_G_ was ∼2-fold) (Fig. 3C). We tried to find mutants that met all three criteria.

The tested 20 AtCnidop3a mutants and their light-dependent cAMP responses are summarized in Fig. 3 and Table 1. Mutants such as T89^2.56^D (Fig. 3D) showed severely reduced light-dependent cellular responses, and mutants such as A292^7.39^D (Fig. 3K) showed similar response patterns to WT (Fig. 3C) (see Table 1). The loss of responses would be due to disruption of protein folding by the mutations, and WT-like responses suggested that the introduced Glu/Asp residues failed to function as the counterion in the resting state. Some mutants such as A90^2.57^E (Fig. 3E), Y113^3.28^D (Fig. 3H), and S186^ECL2^D (Fig. 3J) showed larger green-light induced cAMP elevation than WT, but the cAMP responses were further enhanced by subsequent UV illumination, suggesting that the retinal Schiff base is partially protonated in these mutants. Other mutants such as L94^2.61^D (Fig. 3F) and L94^2.61^E (Fig. 3G) showed robust increases in cAMP levels upon green light illumination, but the responses were not canceled by orange light illumination (L_O_ values are much more than 3), suggesting that absorption spectra of resting and activated states are overlapped and/or bistability was disrupted in these mutants. Finally, one mutant Y113^3.28^E showed a robust and sustained cAMP response upon green light illumination (L_G_ was ∼23-fold), which was comparable to responses after the second UV light illumination (Fig. 3I). The cAMP responses can be turned off by subsequent orange light (L_O_ was ∼2-fold) (Fig. 3I). Taken together, we concluded that AtCniop3a Y113^3.28^E mutant meets all three criteria as visible light sensitive, Gs-coupled, and bistable opsin. Also, the introduced Glu113^3.28^ serves as the counterion for the protonated Schiff base in the resting state of AtCnidop3a.

### Enhancement of green light-induced cellular response of Y113^3.28^E mutant by additional L94^2.61^G substitution

As mentioned above, by introduction of Y113^3.28^E, we successfully converted AtCnidop3a to visible light-sensitive bistable pigment that can be activated and deactivated by green and orange light illuminations, respectively. Although the Y113^3.28^E mutant met our three criteria, the green light-induced cAMP elevation (L_G_ of Y113^3.28^E) was smaller than the UV-induced elevation by WT (L_UV_ of WT) (Fig. 3, C and I, Table 1). We speculated the Y113^3.28^E substitution caused some structural perturbations on arrangement of nearby amino acid residues, leading to negative effect on the opsin’s functionality. We also expected that an additional substitution onto the Y113^3.28^E mutant could compensate the perturbation. In the AlphaFold2 predicted structure of AtCnidop3a, Ala90^2.57^ and Leu94^2.61^ are located near the residue at position 113^3.28^ (Fig. 4A). The additional substitutions at position 90^2.57^ or 94^2.61^ were introduced, and photoresponses of the double mutants were monitored using cAMP GloSensor assay. In particular, we aimed to find double mutants that showed robust green light-induced activation [larger value of L_G-max_ / L_UV_ of WT], sustained cAMP elevation (larger value of L_G-20min_ / L_G-max_), and efficient off response by orange light (L_O_) (Fig. 4B and Table 2). L_G-max_ indicates the maximal response after green light illumination, and L_G-max_ indicates the cAMP response at 20 min after green light illumination (Fig. 4B). In the case of the Y113^3.28^E mutant, the “L_G-max_ / L_UV_ of WT” value was 0.42 and the “L_G-20min_ / L_G-max_” value was 0.56 (Table 2). We tried to find mutants having these values larger than the Y113^3.28^E single mutant

**Fig. 4.**
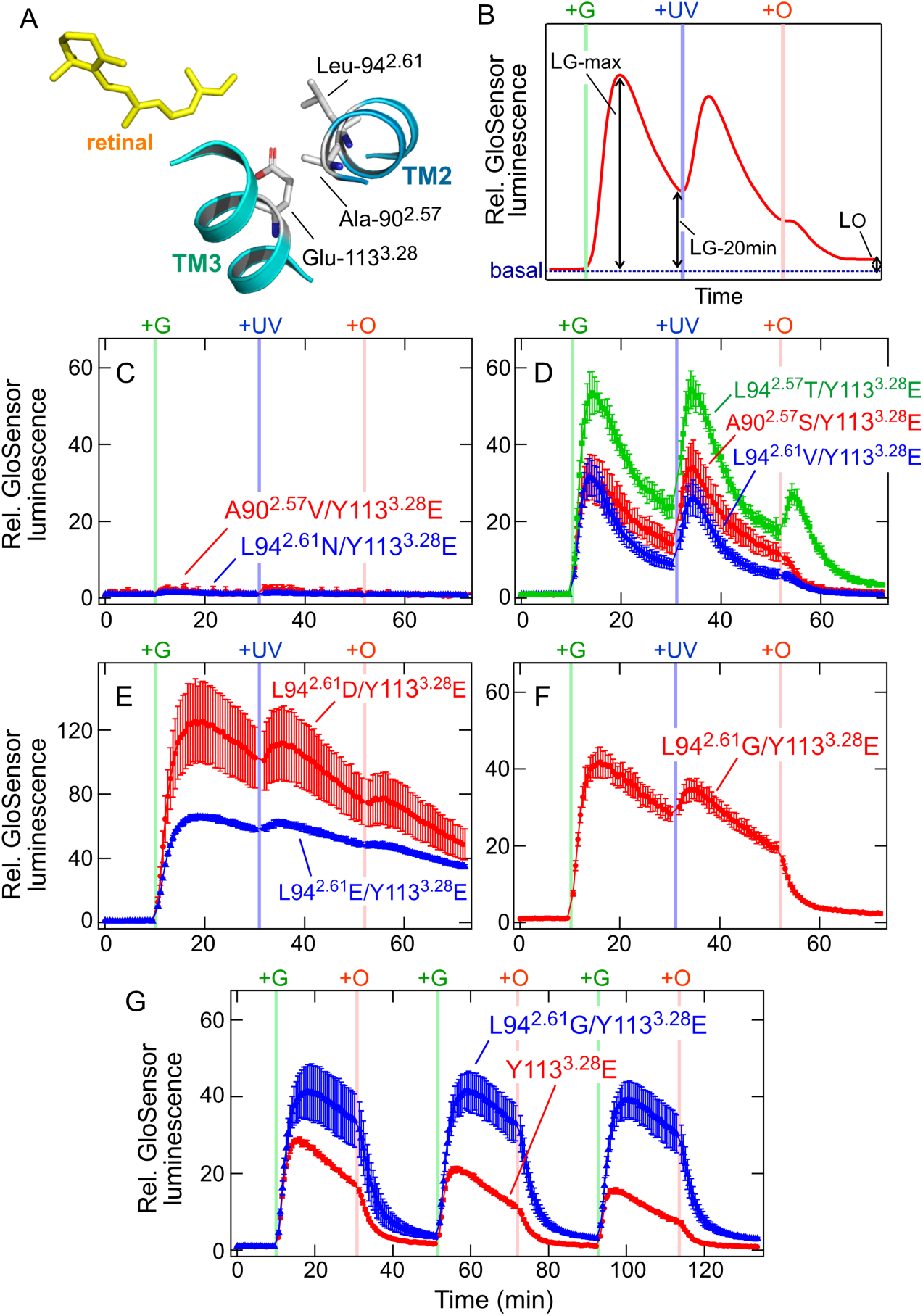
Functional enhancement of visible light-sensitive AtCnidop3a Y113^3.28^E mutant by additional amino acid substitutions. A, Arrangement of the amino acid residues at which we introduced amino acid substitution in this study in the AlphaFold2-predicted structure of AtCnidop3a Y113^3.28^E mutant^41^. The 11-*cis*-retinal molecule is adopted from the crystal structure of bovine rhodopsin (PDB ID: 1U19)^58^. The structural models are prepared using PyMOL (https://pymol.org/). B, Schematic representation of the parameters of GloSensor signals for selection of AtCnidop3a double mutants enhancing the properties of the Y113^3.28^E mutant. “L_G-max_”: the maximal luminescence changes upon the first green light illumination. “L_G-20min_”: the luminescence changes 20 min after the first green light illumination. “L_O_”: the luminescence levels after the third orange-light illumination. C – F, Light-induced changes in cAMP biosensor (GloSensor) luminescence of COS-1 cells expressing AtCnidop3a A90^2.57^V/Y113^3.28^E (*red*) and L94^2.61^N/Y113^3.28^E (*blue*) (C), A90^2.57^S/Y113^3.28^E (*red*), L94^2.61^T/Y113^3.28^E (*green*), and L94^2.61^V/Y113^3.28^E (*blue*) (D), L94^2.61^D/Y113^3.28^E (*red*) and L94^2.61^E/Y113^3.28^E (*blue*) (E), and L94^2.61^G/Y113^3.28^E (F). Luminescence levels are normalized to the value at the starting point (time = 0 min). Green (“G”), UV (“+UV”) and orange (“+O”) bars indicate Green, UV, and orange illuminations, respectively. Error bars indicate the SD values (n= 3). Summary of photoresponses of all tested AtCnidop3a double mutants are shown in Table 2. G, Repetitive increases and decreases in cAMP biosensor (GloSensor) luminescence of COS-1 cells expressing AtCnidop3a mutants Y113^3.28^E (*red*) and L94^2.61^G/Y113^3.28^E (*blue*). Luminescence levels are normalized to the value at the starting point (time = 0 min). Green (“+G”) and orange (“+O”) bars indicate green and orange illuminations, respectively. Error bars indicate the SD values (n= 3 for the Y113^3.28^E mutant and n = 6 for the L94^2.61^G/Y113^3.28^E mutant).

**Table 2.** Light-dependent GloSensor responses of AtCnidop3a double mutants.

| Mutant | LG-max | LG-20min | Lo | LG-max /<br>L<sub>UV</sub> of<br>WT | LG-20min /<br>LG-max |
| --- | --- | --- | --- | --- | --- |
| Y113<sup>3.28</sup>E | 23.4±4.36 | 13.1±2.21 | 1.73±0.16 | 0.42 | 0.56 |
| A90<sup>2.57</sup>C/Y113<sup>3.28</sup>E | 10.8±0.59 | 1.32±0.11 | 0.88±0.16 | 0.19 | 0.12 |
| A90<sup>2.57</sup>D/Y113<sup>3.28</sup>E | 15.8±2.68 | 9.03±1.58 | 3.25±0.30 | 0.29 | 0.57 |
| A90<sup>2.57</sup>E/Y113<sup>3.28</sup>E | 11.8±1.44 | 7.22±0.66 | 3.61±0.36 | 0.21 | 0.61 |
| A90<sup>2.57</sup>F/Y113<sup>3.28</sup>E | 5.22±1.02 | 1.90±0.30 | 0.58±0.17 | 0.09 | 0.36 |
| A90<sup>2.57</sup>G/Y113<sup>3.28</sup>E | 22.3±6.61 | 10.1±3.98 | 1.00±0.16 | 0.40 | 0.45 |
| A90<sup>2.57</sup>H/Y113<sup>3.28</sup>E | 6.10±1.34 | 1.14±0.41 | 0.57±0.21 | 0.11 | 0.19 |
| A90<sup>2.57</sup>I/Y113<sup>3.28</sup>E | 2.49±0.63 | 1.14±0.34 | 0.51±0.13 | 0.05 | 0.46 |
| A90<sup>2.57</sup>K/Y113<sup>3.28</sup>E | 1.93±0.22 | 1.51±0.45 | 0.86±0.37 | 0.04 | 0.78 |
| A90<sup>2.57</sup>L/Y113<sup>3.28</sup>E | 4.19±1.17 | 1.04±0.15 | 0.58±0.14 | 0.08 | 0.25 |
| A90<sup>2.57</sup>M/Y113<sup>3.28</sup>E | 17.5±1.08 | 2.90±0.33 | 1.20±0.16 | 0.32 | 0.17 |
| A90<sup>2.57</sup>N/Y113<sup>3.28</sup>E | 14.4±2.44 | 6.46±0.93 | 2.60±0.48 | 0.26 | 0.45 |
| A90<sup>2.57</sup>P/Y113<sup>3.28</sup>E | 3.23±0.16 | 1.93±0.36 | 1.07±0.17 | 0.06 | 0.60 |
| A90<sup>2.57</sup>Q/Y113<sup>3.28</sup>E | 25.1±1.27 | 10.9±0.49 | 1.78±0.15 | 0.46 | 0.43 |
| A90<sup>2.57</sup>R/Y113<sup>3.28</sup>E | 6.01±1.17 | 3.18±0.33 | 1.66±0.31 | 0.11 | 0.53 |
| A90<sup>2.57</sup>S/Y113<sup>3.28</sup>E | 32.2±5.29 | 14.2±2.67 | 1.45±0.12 | 0.58 | 0.44 |
| A90<sup>2.57</sup>T/Y113<sup>3.28</sup>E | 9.47±1.63 | 1.95±0.36 | 1.02±0.14 | 0.17 | 0.21 |
| A90<sup>2.57</sup>V/Y113<sup>3.28</sup>E | 2.63±0.90 | 0.96±0.11 | 0.52±0.10 | 0.05 | 0.37 |
| A90<sup>2.57</sup>W/Y113<sup>3.28</sup>E | 1.75±0.19 | 1.01±0.12 | 0.61±0.29 | 0.03 | 0.58 |
| A90<sup>2.57</sup>Y/Y113<sup>3.28</sup>E | 1.06±0.05 | 0.90±0.03 | 0.32±0.03 | 0.02 | 0.85 |
| L94<sup>2.61</sup>A/Y113<sup>3.28</sup>E | 29.9±8.31 | 13.8±3.93 | 1.62±0.12 | 0.54 | 0.46 |
| L94<sup>2.61</sup>C/Y113<sup>3.28</sup>E | 28.9±1.20 | 9.03±0.45 | 1.08±0.09 | 0.52 | 0.31 |
| L94<sup>2.61</sup>D/Y113<sup>3.28</sup>E | 126±26.5 | 103±19.0 | 48.8±9.63 | 2.29 | 0.82 |
| L94<sup>2.61</sup>E/Y113<sup>3.28</sup>E | 66.0±1.45 | 58.2±1.28 | 34.9±1.40 | 1.20 | 0.88 |
| L94<sup>2.61</sup>F/Y113<sup>3.28</sup>E | 8.80±0.37 | 5.65±0.54 | 1.17±0.12 | 0.16 | 0.64 |
| L94<sup>2.61</sup>G/Y113<sup>3.28</sup>E | 41.8±3.95 | 28.3±2.92 | 2.30±0.10 | 0.76 | 0.68 |
| L94<sup>2.61</sup>H/Y113<sup>3.28</sup>E | 20.9±5.38 | 3.71±0.72 | 0.99±0.18 | 0.38 | 0.18 |
| L94<sup>2.61</sup>I/Y113<sup>3.28</sup>E | 18.1±1.36 | 6.99±0.40 | 1.09±0.09 | 0.33 | 0.39 |
| L94<sup>2.61</sup>K/Y113<sup>3.28</sup>E | 1.77±0.25 | 1.15±0.16 | 0.80±0.06 | 0.03 | 0.65 |
| L94<sup>2.61</sup>M/Y113<sup>3.28</sup>E | 16.2±0.85 | 6.06±0.28 | 1.18±0.05 | 0.29 | 0.37 |
| L94<sup>2.61</sup>N/Y113<sup>3.28</sup>E | 1.60±0.20 | 1.06±0.10 | 0.73±0.05 | 0.03 | 0.66 |
| L94<sup>2.61</sup>P/Y113<sup>3.28</sup>E | 2.04±0.05 | 1.31±0.05 | 0.96±0.12 | 0.04 | 0.64 |
| L94<sup>2.61</sup>Q/Y113<sup>3.28</sup>E | 22.1±0.95 | 8.55±0.33 | 1.53±0.04 | 0.40 | 0.39 |
| L94<sup>2.61</sup>R/Y113<sup>3.28</sup>E | 12.8±0.70 | 8.19±0.52 | 2.68±0.07 | 0.23 | 0.64 |
| L94<sup>2.61</sup>S/Y113<sup>3.28</sup>E | 43.6±9.52 | 18.1±3.61 | 2.73±0.51 | 0.79 | 0.42 |
| L94<sup>2.61</sup>T/Y113<sup>3.28</sup>E | 53.5±5.60 | 23.8±2.88 | 3.46±0.22 | 0.97 | 0.44 |
| L94<sup>2.61</sup>V/Y113<sup>3.28</sup>E | 31.5±4.89 | 8.78±1.29 | 1.19±0.26 | 0.57 | 0.28 |
| L94<sup>2.61</sup>W/Y113<sup>3.28</sup>E | 3.73±0.41 | 2.36±0.14 | 0.99±0.04 | 0.07 | 0.63 |
| L94<sup>2.61</sup>Y/Y113<sup>3.28</sup>E | 12.1±0.92 | 3.54±0.26 | 1.21±0.10 | 0.22 | 0.29 |

Tested double mutants and their light-dependent cAMP responses are summarized in Fig. 4 and Table 2. Among 38 double mutants, double mutants such as A90^2.57^V/Y113^3.28^E and L94^2.61^N/Y113^3.28^E showed little cAMP responses upon light illuminations (Fig. 4C), probably due to disruption of protein folding by the additional substitutions. Some double mutants such as A90^2.57^S/Y113^3.28^E, L94^2.61^T/Y113^3.28^E, and L94^2.61^V/Y113^3.28^E evoked cAMP elevation upon green-light illumination, but the responses were transient (“L_G-20min_ / L_G-max_” values were less than 0.5) (Fig. 4D). In the case of the L94^2.61^T/Y113^3.28^E mutant, even the third orange-light illumination caused a small and transient cAMP response (Fig. 4D), suggesting that the resting state can absorb orange light. Mutants L94^2.61^D/Y113^3.28^E and L94^2.61^E/Y113^3.28^E showed larger green light-induced cAMP responses than UV-induced WT response (Fig. 4E), but the responses were not canceled by orange light (large L_O_ values) like the cases of L94^2.61^D and L94^2.61^E single mutants (Fig. 3, F and G). These results suggested that the mutational effects by L94^2.61^D or L94^2.61^E substitution are superior to the effect of Y113^3.28^E substitution. We finally selected the L94^2.61^G/Y113^3.28^E double mutant, because the mutant showed a larger cAMP elevation upon green-light illumination than the Y113^3.28^E mutant, the cAMP response was sustained, and the response was canceled by orange-light illumination (Fig. 4F). Both the “L_G-max_ / L_UV_ of WT” value (0.76) and the “L_G-20min_ / L_G-max_” value (0.68) of the double mutant were larger than those of the Y113^3.28^E single mutant (Table 2). These results suggested that the additional L94^2.61^G substitution on Y113^3.28^E mutant does not seriously affect activation and deactivation processes in the opsin, but the additional substitution reduces some structural perturbations in the Y113^3.28^E single mutant. The green light-induced activation and the orange light-induced deactivation of the L94^2.61^G/Y113^3.28^E mutant as well as the Y113^3.28^E mutant can be repeated (Fig. 4G).

In order to reveal the green light-sensitive spectral properties of the L94^2.61^G/Y113^3.28^E mutant, we expressed in COS-1 cells, extracted and purified the double mutant. We expected that purified L94^2.61^G/Y113^3.28^E mutants showed somewhat absorbance around 520 nm in the visible region. Unfortunately, the obtained “purified” sample of the mutant did not show absorbance at UV and visible region other than 280-nm protein absorbance (Fig. 5A). This failure of the pigment purification is probably because the substitutions L94^2.61^G/Y113^3.28^E negatively affected expression levels in the cultured cell and/or stability of the protein folding under the solubilized conditions.

**Fig. 5.**
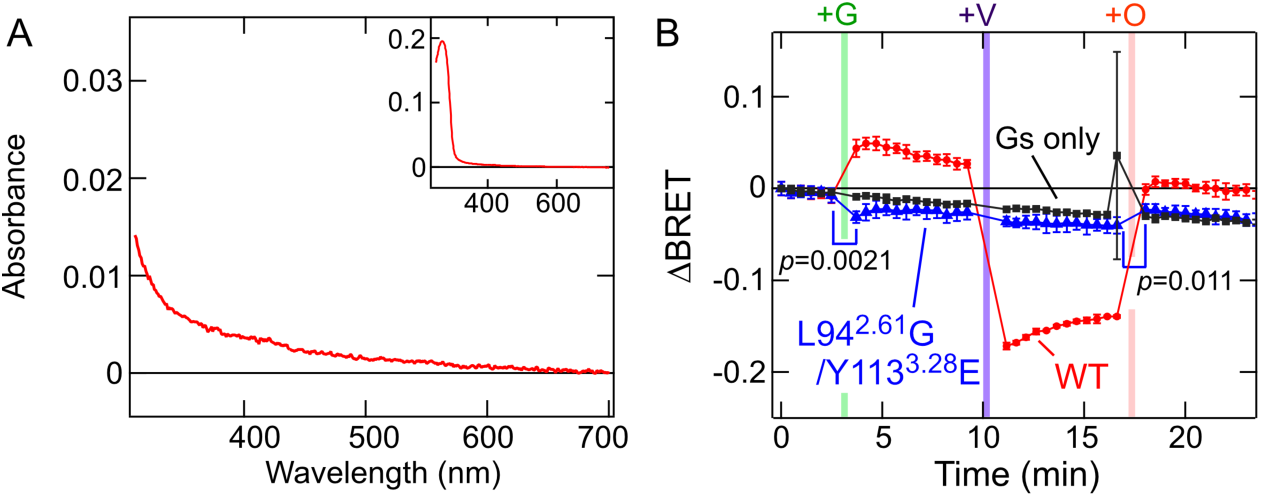
Spectral properties and Gs activation/deactivation of AtCnidop3a L94^2.61^G/Y113^3.28^E double mutant. A, Absorption spectrum of purified AtCnidop3a L94^2.61^G/Y113^3.28^E mutant in the dark (*red*) is shown. (*Inset*) Absorption spectrum of the L94^2.61^G/Y113^3.28^E mutant in a wider wavelength range. B, Light-induced changes in the ΔBRET values of luminescence of HEK293T Gs/Gq/Gi KO cells expressing AtCnidop3a WT (*red*), L94^2.61^G/Y113^3.28^E mutant (blue), and without the opsin (*black*). ΔBRET time courses upon green- (“+G”), violet- (“+V”) and orange- (“+O”) light illuminations are shown. Error bars indicate the SD values (n= 3). The green and orange light-induced ΔBRET changes are statistically significant (p values are 0.0021 and 0.011, respectively) based on paired Student’s t-test.

We next tried to prove the green light-sensitivity of the L94^2.61^G/Y113^3.28^E mutant using the BRET-based Gs dissociation assay. In Fig. 5B, we showed green light-induced Gs trimer dissociation and orange light-induced (re)association. In comparison with UV-induced Gs dissociation signals by WT (see also Figs. 2B), the L94^2.61^G/Y113^3.28^E mutant showed much smaller dissociation signals upon green light illumination. The dissociation BRET signals were cancelled by orange light illumination (Fig. 5B). The green and orange light-induced BRET changes of the L94^2.61^G/Y113^3.28^E mutant are small but significant (Fig. 5B). These results suggest that expression levels of the L94^2.61^G/Y113^3.28^E in mammalian cultured cells were much decreased by the double substitutions. On the other hand, cAMP responses would be amplified through signal transmission if the opsin, Gsα, and adenylyl cyclase, leading to large cAMP responses (but less than that of WT) (Figs. 3C and 4F).

Our spectral and cellular assays on AtCnidop3a revealed that the opsin is UV-sensitive Gs-coupled bistable opsin that can be deactivated by orange light. The opsin can bidirectionally modulate intracellular cAMP levels by UV and orange light stimulations. Furthermore, the substitution of the L94^2.61^G/Y113^3.28^E in the opsin converted it to be activated by green light while retaining the property of being deactivated by orange light. The insights of this study will help understanding of UV-dependent physiological responses of the coral and engineering of a novel optical control tool modulating Gs-dependent cellular responses (see “Discussion”).

## Discussion

In this study, we revealed molecular characteristics of AtCnidop3a and engineered the functional properties via site-directed mutagenesis. In this section, we discuss relationship between the AtCnidop3a properties and light responses of the coral species. We also discuss how spectral properties of cnidopsins have been tuned, and discuss utility of AtCnidop3a WT and L94^2.61^G/Y113^3.28^E mutant as optical control tools to regulate intercellular signalings upon different color of lights.

### Molecular properties of AtCnidopsin3 and UV-dependent physiological responses of the coral *Acropora tenuis*

Our data clearly indicated that AtCnidop3a is a UV-sensitive, Gs-coupled, and bistable opsin. The bistable property enables the opsin to detect relative intensity of environmental light around 400 nm (UV to violet region) and around 550 nm (green to yellow region). A previous study reported that larvae of *Acropora tenuis* stop to swim upon attenuation of short-wavelength light^3^. AtCnidop3a could play an important role in regulation of the swimming behavior. In addition, previous studies showed that another cnidopsin (AtCnidop7) and a different-type opsin Antho2e, belonging to a phylogenetically different anthozoan-specific opsin II (ASO-II) group, are also UV-sensitive pigments^16,18^. AtCnidop7 would induce cAMP responses via Gs activation like other cnidopsins, and Antho2e might drive Ca^2+^ signalings like Antho2a belonging to the ASO-II group. These results and insights suggested that UV-sensing in *Acropora tenuis* is mediated by multiple opsins including AtCnidop3a to produce complicated cellular responses. Further studies using the coral organisms applying genetic modification approach such as CRISPR/Cas9 based gene knockout^48–50^ are needed to reveal which opsins (or other photoreceptive molecules) are responsible for UV-dependent photoresponses in *Acropora tenuis*.

### Spectral tuning via protonation and deprotonation at the retinal Schiff base in cnidopsins

As mentioned in “Results”, AtCnidop3a WT is UV-sensitive and introduction of a Glu or Asp residue at some positions such as position 113^3.28^ made the opsin be activated by visible (green) light (Fig. 3 and Table 1). Thus, in the resting state of WT, the retinal Schiff base would be deprotonated due to lack of the counterion. Previous study also reported that in the resting state of JellyOP, the introduced Glu residue at position 113^3.28^ acts as the counterion^35^. Furthermore, Glu113^3.28^ functions as a natural counterion for retinal Schiff base in vertebrate visual pigment such as bovine rhodopsin^44,51,52^. These results suggested that arrangement of the amino acid residue at position 113^3.28^ and the Schiff base is conserved among the resting state of various opsins. In contrast, AtCnidop3a is bistable, but JellyOP and vertebrate visual pigments are unidirectionally activated by light^12,53^, suggesting that the arrangement is somewhat diversified in the activated states of opsins, as discussed in a previous paper^40^.

A recent study reported that in another cnidarian opsin Antho2a in the ASO-II group, chloride ion rather than specific amino acid residue acts as the counterion^16^. Chloride (and other negative charged) ions could participate in stabilization of the proton on the retinal Schiff base in cnidopsins. Further analysis is needed to understand spectral tuning mechanism involving deprotonation or protonation of retinal Schiff base in cnidopsins.

### Potentials of AtCnidop3a WT and L94^2.61^G/Y113^3.28^E mutant as useful optical control tools

Since the discovery of Gs-coupled JellyOP^12^, the cnidopsin has been used as a useful optogenetic tool to modulate intracellular cAMP responses^19,20,36^. Intracellular cAMP elevation can cause various cellular responses such as activation of PKA and modulation of CNG and HCN channels^54^. Optogenetic manipulation of cAMP levels using Gs-independent tools are also reported^55,56^. As an optogenetic tool, JellyOP induces transient cAMP elevation upon green-light illumination followed by spontaneous decay of the signal^36^. AtCnidopsin3a could be also used as another valuable optogenetic tool modulating cAMP responses. In comparison with JellyOP, AtCnidop3a WT can induce more sustained cAMP responses by UV-light illumination, and the responses can be turned off by orange-light illumination (Fig. 2D). If researchers want to use visible light for not only deactivation but activation, AtCnidop3a L94^2.61^G/Y113^3.28^E mutant would be suitable for the purpose (Fig. 4G). Taken together, AtCnidop3a WT and L94^2.61^G/Y113^3.28^E mutant can be valuable optical control tools to achieve repeatable and switchable manipulation of Gs-dependent intercellular cAMP responses by stimulation with different wavelengths of light.

## Acknowledgement

We thank Dr. David Farrens (Oregon Health & Science University) for providing us with COS-1 cell line and the pMT expression vector, Dr. Robert Molday (University of British Columbia, Canada) for providing hybridoma cells producing 1D4 antibody, and Dr. Asuka Inoue (Kyoto University, Japan) for providing the RIC8B plasmid. We also thank Dr. Akihisa Terakita and Dr. Mitsumasa Koyanagi (Osaka Metropolitan University, Japan) for valuable advices to set up the GloSensor assay system for AtCnidop3a, and Dr. Kunio Inoue, Dr. Saori Tani-Matsuhana, Yuri Asai, Naoyuki Taira, and Yusei Sakata (Kobe University, Japan) for discussing the potential of AtCnidop3a as optogenetic tools on zebrafish.

## Author contribution

Y. S., A. S., and H. T. designed the study, conducted experiments, and analyzed obtained data. T. K. and K. T. set up and conducted the BRET-based experiments. K. S., K. K., and H. O. established the Gs/Gq/Gi KO HEK293T cells and tested experimental conditions using the KO cells. All authors discussed obtained data and wrote the paper.

## Abbreviations

BRET: bioluminescence resonance energy transfer
Gs: Gs-type trimeric G proteins
GPCR: G protein-coupled receptor
λmax: absorption maximum
ML: maximum likelihood
UV: ultra-violet
WT: wild-type

