## Supplemental Information for "A bistable UV-sensitive opsin from a reef building coral showing a switchable and tunable regulation of Gs-signaling by different wavelengths of light"

**Supplemental Figs. S1, S2, S3, and S4 with their captions**
**Supplemental Tables S1 and S2**
**Supplemental Data**

Yusuke Sakai*#[1, 2], Akinari Sakayori#[3], Tomoki Kawaguchi[3], Kento Takano[3], Keita Sato[4], Keiichi Kojima[4], Hideyo Ohuchi[4], Hisao Tsukamoto*[3].

[1]Division of Neuroscience and Centre for Biological Timing, School of Biological Sciences, Faculty of Biology Medicine and Health, University of Manchester, Manchester, UK; [2]Department of Biology, Graduate School of Science, Osaka Metropolitan University, Osaka, Japan; [3]Department of Biology, Graduate School of Science, Kobe University, Kobe, Japan; [4]Faculty of Medicine, Dentistry, and Pharmaceutical Sciences, Okayama University, Okayama, Japan.

#These authors equally contributed to this study.

*Correspondence should be addressed to Hisao Tsukamoto; Department of Biology, Graduate School of Science, Kobe University, 1-1, Rokkodai-cho, Nada-Ku, Kobe, 657-8501, Japan;, or Yusuke Sakai, Department of Biology, Graduate School of Science, Osaka Metropolitan University, 3-3-138, Sugimoto-cho, Sumiyorhi-Ku, Osaka, 558-8585, Japan;

### Supplemental figures

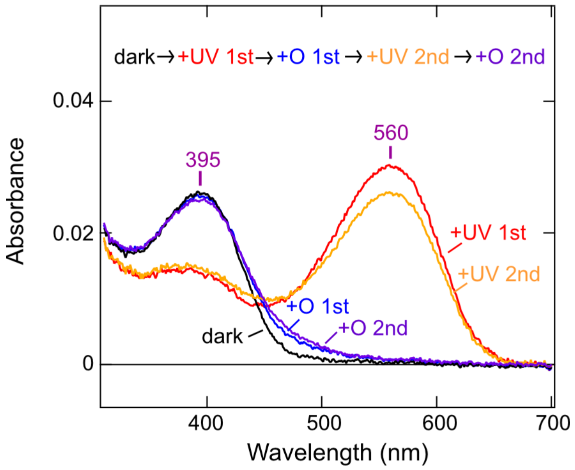


**<u>Supplemental Fig. S1</u> Spectral changes of AtCnidop3a upon repeated UV-orange light illuminations.**

Absorption spectra of purified AtCnidop3a WT in the dark (black), after 1st UV illumination (red), after 1st orange light illumination (blue), after 2nd UV illumination (orange), and 2nd orange light illumination (violet) are shown. The order of light illuminations is also indicated.

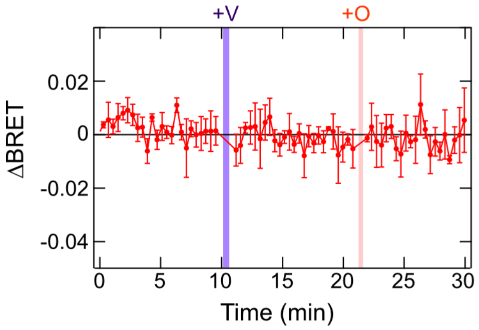


**<u>Supplemental Fig. S2</u> BRET-based Gs dissociation assay on AtCnidop3a using COS-1 cells.**

ΔBRET time course of COS-1 cells expressing AtCnidop3a WT (*red*) upon violet- (“+V”) and orange- (“+O”) light illuminations is shown. Error bars indicate the SD values (n= 3).

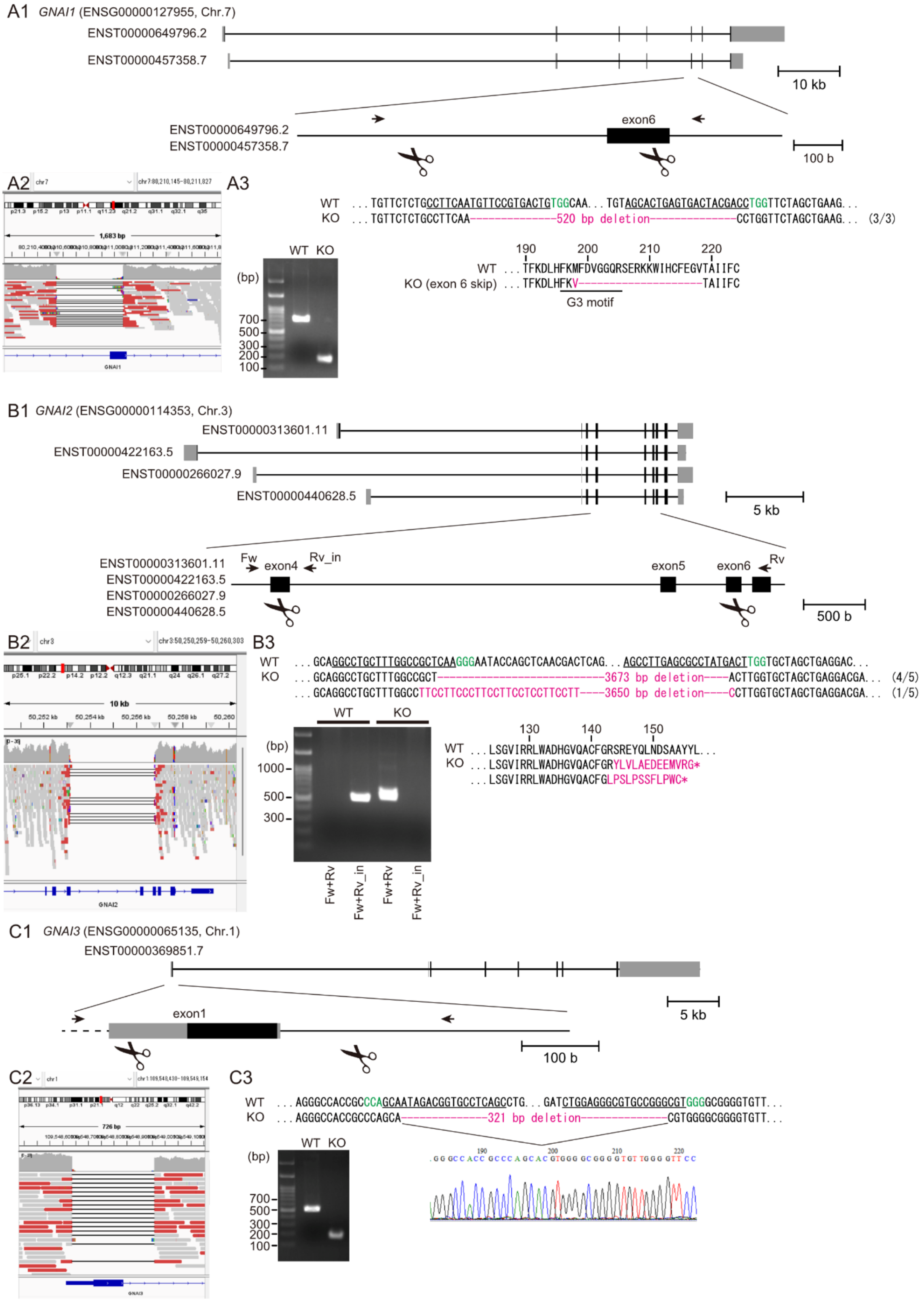

A1 GNAI1 (ENSG00000127955, Chr.7)
ENST00000649796.2
ENST00000457358.7
10 kb
exon6
100 b
A2
A3
WT ...TGTTCTCTGCCTTCAATGTTCCGTGACTGTGGCAA...TGTAGCACTGAGTGACTACGACCTGGTTCTAGCTGAAG...
KO ...TGTTCTCTGCCTTCAA-------------520 bp deletion-------------CCTGGTTCTAGCTGAAG... (3/3)
WT KO
(bp)
700
500
300
200
100
190 200 210 220
WT ...TFKDLHFKMFDVGGQRSERKKWIHCFEGVTAIIFC
KO (exon 6 skip) ...TFKDLHFKV------------------TAIIFC
G3 motif
B1 GNAI2 (ENSG00000114353, Chr.3)
ENST00000313601.11
ENST00000422163.5
ENST00000266027.9
ENST00000440628.5
5 kb
Fw
exon4
Rv_in
exon5
exon6
Rv
500 b
B2
B3
WT ...GCAGGCCTGCTTTGGCCGCTCAAGGGAATACCAGCTCAACGACTCAG...AGCCTTGAGCGCCTATGACTTGGTGCTAGCTGAGGAC...
KO ...GCAGGCCTGCTTTGGCCGCT------------------------3673 bp deletion----ACTTGGTGCTAGCTGAGGACGA... (4/5)
...GCAGGCCTGCTTTGGCCTTCCTTCCCTTCCTTCCTCCTTCCTT----3650 bp deletion----CCTTGGTGCTAGCTGAGGACGA... (1/5)
WT KO
(bp)
1000
500
300
Fw+Rv
Fw+Rv_in
Fw+Rv
Fw+Rv_in
130 140 150
WT ...LSGVIRRLWADHGVQACFGRSREYQLNDSAAYYL...
KO ...LSGVIRRLWADHGVQACFGRYLVLAEDEEMVRG*
...LSGVIRRLWADHGVQACFGLPSLPSSFLPWC*
C1 GNAI3 (ENSG00000065135, Chr.1)
ENST00000369851.7
5 kb
exon1
100 b
C2
C3
WT ...AGGGCCACCGCCCAGCAATAGACGGTGCCTCAGCCTG...GATCTGGAGGGCGTGCCGGGCGTGGGGCGGGGTGTT...
KO ...AGGGCCACCGCCCAGCA-------------321 bp deletion-------------CGTGGGGCGGGGTGTT...
WT KO
(bp)
700
500
300
200
100
190 200 210 220

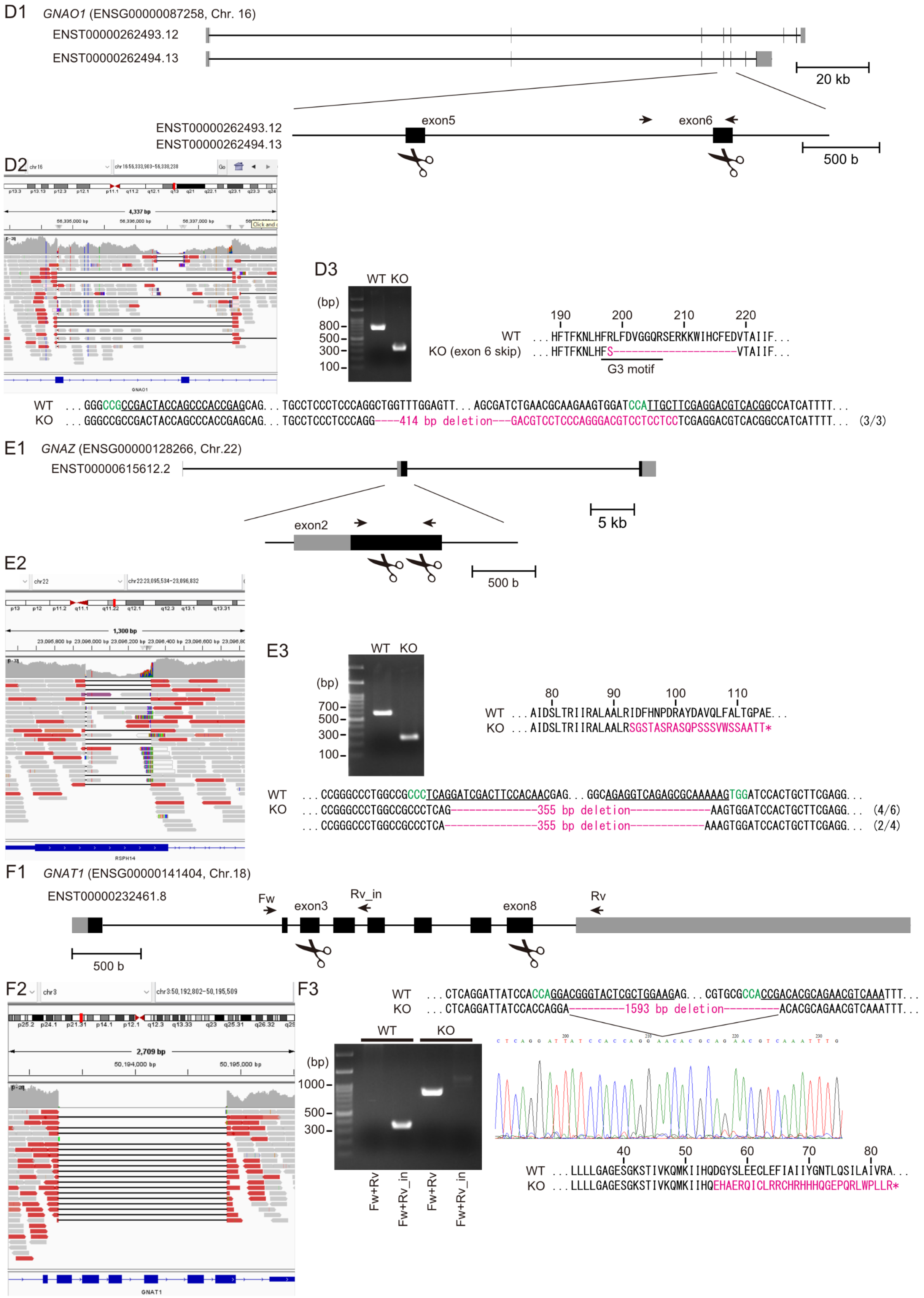
D1 GNAO1 (ENSG00000087258, Chr. 16)
ENST00000262493.12
ENST00000262494.13
20 kb
exon5
exon6
500 b
D2
D3
WT KO
(bp)
800
500
300
100
190 200 210 220
WT ...HFTFKNLHFRLFDVGGQRSERKKWIHCFEDVTAIIF...
KO (exon 6 skip) ...HFTFKNLHFS--------------------VTAIIF...
G3 motif
WT ...GGGCCGCCGACTACCAGCCCACCGAGCAG...TGCCTCCCTCCCAGGCTGGTTTGGAGTT...AGCGATCTGAACGCAAGAAGTGGATCCATTGCTTCGAGGACGTCACGGCCATCATTTT...
KO ...GGGCCGCCGACTACCAGCCCACCGAGCAG...TGCCTCCCTCCCAGG----414 bp deletion---GACGTCCTCCCAGGGACGTCCTCCTCCTCGAGGACGTCACGGCCATCATTTT... (3/3)
E1 GNAZ (ENSG00000128266, Chr.22)
ENST00000615612.2
5 kb
exon2
500 b
E2
E3
WT KO
(bp)
700
500
300
100
80 90 100 110
WT ...AIDSLTRIIRALAALRIDFHNPDRAYDAVQLFALTGPAE...
KO ...AIDSLTRIIRALAALRSGSTASRASQPSSSVWSSAATT*
WT ...CCGGGCCCTGGCCGCCCTCAGGATCGACTTCCACAACGAG...GGCAGAGGTCAGAGCGCAAAAAGTGGATCCACTGCTTCGAGG...
KO ...CCGGGCCCTGGCCGCCCTCAG---------------355 bp deletion--------------AAGTGGATCCACTGCTTCGAGG... (4/6)
...CCGGGCCCTGGCCGCCCTCA----------------355 bp deletion-------------AAAGTGGATCCACTGCTTCGAGG... (2/4)
F1 GNAT1 (ENSG00000141404, Chr.18)
ENST00000232461.8
Fw
exon3
Rv_in
exon8
Rv
500 b
F2
F3
WT ...CTCAGGATTATCCACCAGGACGGGTACTCGCTGGAAGAG...CGTGCGCCACCGACACGCAGAACGTCAAATTT...
KO ...CTCAGGATTATCCACCAGGA---------1593 bp deletion---------ACACGCAGAACGTCAAATTT...
WT KO
(bp)
1000
500
300
Fw+Rv
Fw+Rv_in
Fw+Rv
Fw+Rv_in
40 50 60 70 80
WT ...LLLLGAGESGKSTIVKQMKIIHQDGYSLEECLEFIAIIYGNTLQSILAIVRA...
KO ...LLLLGAGESGKSTIVKQMKIIHQEHAERQICLRRCHRHHHQGEPQRLWPLLR*

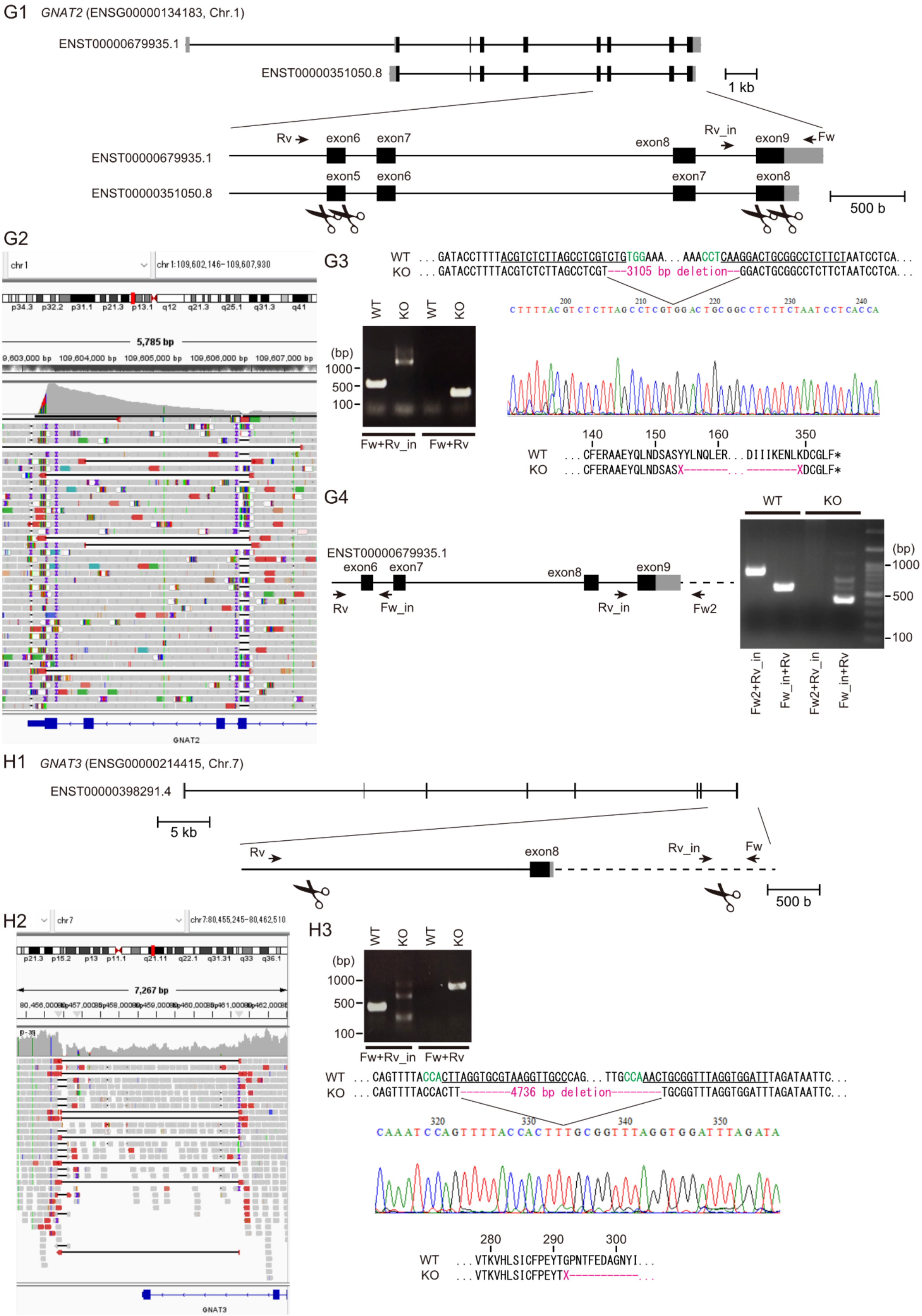
G1 GNAT2 (ENSG00000134183, Chr.1)
ENST00000679935.1
ENST00000351050.8
1 kb
Rv
exon6
exon7
exon8
Rv_in
exon9
Fw
exon5
exon6
exon7
exon8
500 b
G2
G3
WT ...GATACCTTTTACGTCTCTTAGCCTCGTCTGTGGAAA...AAACCTCAAGGACTGCGGCCTCTTCTAATCCTCA...
KO ...GATACCTTTTACGTCTCTTAGCCTCGT---3105 bp deletion--GGACTGCGGCCTCTTCTAATCCTCA...
WT
KO
(bp)
1000
500
100
Fw+Rv_in
Fw+Rv
140 150 160 350
WT ...CFERAAEYQLNDSASYYLNQLER...DIIKENLKDCGLF*
KO ...CFERAAEYQLNDSASX-------...--------XDCGLF*
G4
ENST00000679935.1
Rv
Fw_in
Rv_in
Fw2
(bp)
1000
500
100
Fw2+Rv_in
Fw_in+Rv
H1 GNAT3 (ENSG00000214415, Chr.7)
ENST00000398291.4
5 kb
exon8
500 b
H2
H3
WT ...CAGTTTTACCACTTAGGTGCGTAAGGTTGCCCAG...TTGCCAAACTGCGGTTTAGGTGGATTTAGATAATTC...
KO ...CAGTTTTACCACTT--------4736 bp deletion--------TGCGGTTTAGGTGGATTTAGATAATTC...
280 290 300
WT ...VTKVHLSICFPEYTGPNTFEDAGNYI...
KO ...VTKVHLSICFPEYTX------------...

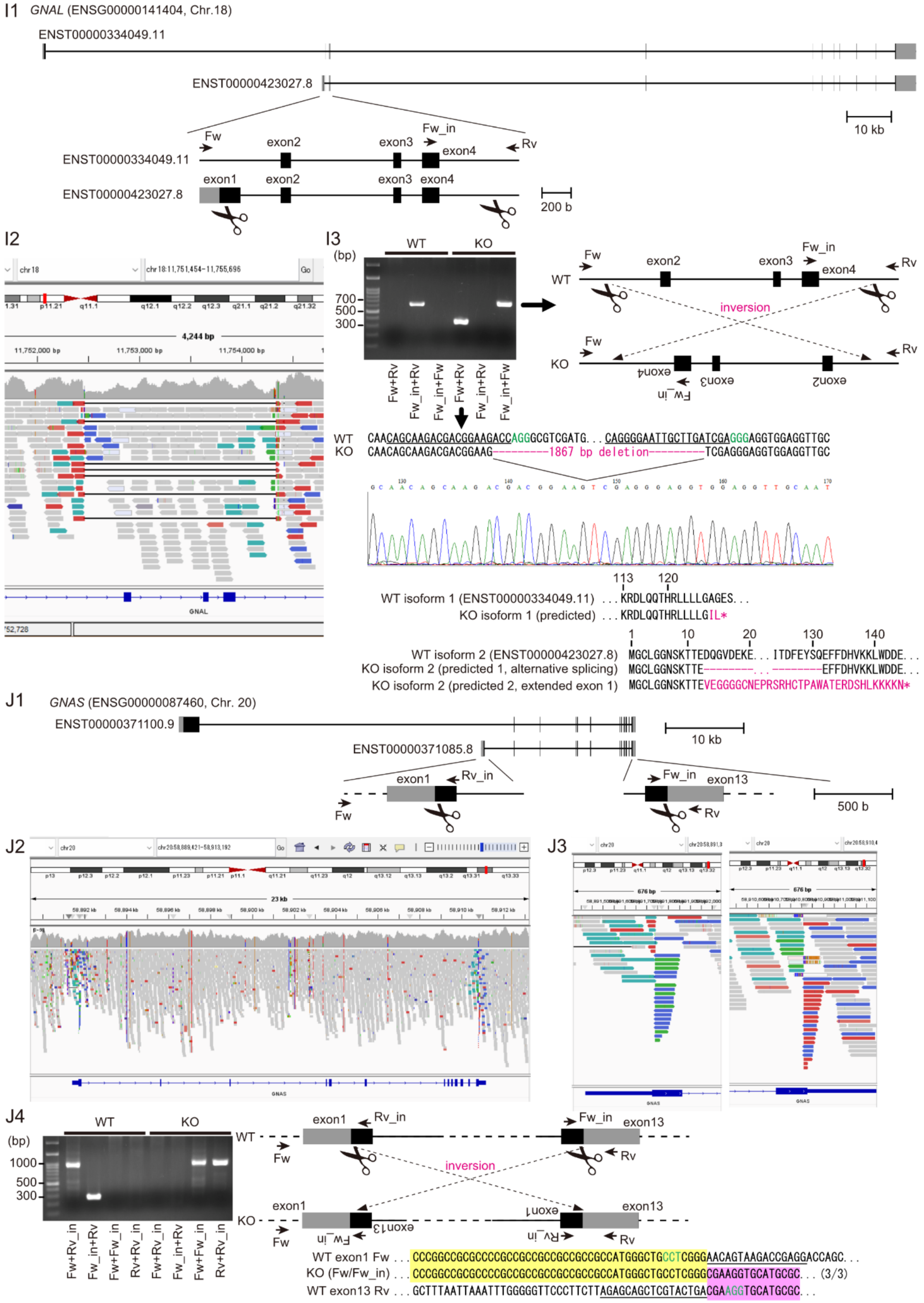
I1
GNAL (ENSG00000141404, Chr.18)
ENST00000334049.11
ENST00000423027.8
10 kb
Fw
exon2
exon3
Fw_in
exon4
Rv
exon1
200 b
I2
I3
(bp)
WT
KO
700
500
300
Fw+Rv
Fw_in+Rv
Fw_in+Fw
inversion
WT CAACAGCAAGACGACGGAAGACCAGGGCGTCGATG. . . CAGGGGAATTGCTTGATCGAGGGAGGTGGAGGTTGC
KO CAACAGCAAGACGACGGAAG---------1867 bp deletion---------TCGAGGGAGGTGGAGGTTGC
113 120
WT isoform 1 (ENST00000334049.11) . . . KRDLQQTHRLLLLGAGES. . .
KO isoform 1 (predicted) . . . KRDLQQTHRLLLLGIL*
1 10 20 130 140
WT isoform 2 (ENST00000423027.8) MGCLGGNSKTTEDQGVDEKE. . . ITDFEYSQEFFDHVKKLWDDE. . .
KO isoform 2 (predicted 1, alternative splicing) MGCLGGNSKTTE--------. . . --------EFFDHVKKLWDDE. . .
KO isoform 2 (predicted 2, extended exon 1) MGCLGGNSKTTEVEGGGGCNEPRSRHCTPAWATERDSHLKKKKN*
J1
GNAS (ENSG00000087460, Chr. 20)
ENST00000371100.9
ENST00000371085.8
10 kb
exon1
Rv_in
Fw_in
exon13
Fw
Rv
500 b
J2
J3
J4
(bp)
1000
500
300
Fw+Rv_in
Fw_in+Rv
Fw+Fw_in
Rv+Rv_in
inversion
WT exon1 Fw . . . CCCGGCCGCGCCCCGCCGCCGCCGCCGCCGCCATGGGCTGCCTCGGGAACAGTAAGACCGAGGACCAGC. . .
KO (Fw/Fw_in) . . . CCCGGCCGCGCCCCGCCGCCGCCGCCGCCGCCATGGGCTGCCTCGGGCGAAGGTGCATGCGC. . . (3/3)
WT exon13 Rv . . . GCTTTAATTAAATTTGGGGGTTCCCTTCTTAGAGCAGCTCGTACTGACGAAGGTGCATGCGC. . .

**<u>Supplemental Fig. S3</u> Target sequences used for genome editing and the resulting genotypes of the Gs/Gi/Gq KO HEK293T cells generated in this study.**

To confirm genome editing, genomic PCR products were directly sequenced for *GNAI3*, *GNAT1*, *GNAT2*, *GNAT3*, and *GNAL*. For *GNAI1*, *GNAI2*, *GNAO1*, *GNAZ*, and *GNAS* genomic PCR products were ligated into the pMD20 T-vector (Takara, Japan), transformed into *E. coli*, and sequenced after colony PCR. In the schematic drawings, black and gray boxes indicate coding and untranslated regions, respectively. Arrows and scissors indicate the positions of primers used for genotyping and the sites targeted for SpCas9 cleavage, respectively. Green letters and underlined sequences in the nucleotide alignments indicate protospacer adjacent motifs (PAMs) and sgRNA target sequences, respectively. Magenta letters in the nucleotide and amino acid sequences indicate deletions or replacements in the mutant alleles. (A1) Gene structure, sgRNA-targeted sites, and primer binding sites of *GNAI1*. (A2) IGV view of the targeted region and surrounding sequences at the *GNAI1* locus in the whole-genome sequencing data. (A3) Genomic PCR and sequences determined from cloned colonies. Three cloned colonies were analyzed, and all contained a 520-bp deletion including exon 6. The predicted amino acid sequence is shown below. Deletion of amino acids F199-V218 removes the essential GTP-binding G3 motif. (B1) Gene structure, sgRNA-targeted sites, and primer binding sites of *GNAI2*. (B2) IGV view of the targeted region and surrounding sequences at the *GNAI2* locus in the whole-genome sequencing data. (B3) Genomic PCR performed with primer pairs Fw/Rv_in and Fw/Rv, and sequences determined from cloned colonies. The binding site for Rv_in is deleted in the KO allele. Because the distance between Fw and Rv in the WT allele was too long to be amplified under these conditions, a band was detected with Fw/Rv only in the KO allele. Among five cloned colonies analyzed, four and one carried 3673-bp and 3650-bp deletions, respectively. The predicted amino acid sequences deduced from the mutant alleles are C-terminally truncated by frameshift. (C1) Gene structure, sgRNA-targeted sites, and primer binding sites of *GNAI3*. (C2) IGV view of the targeted region and surrounding sequences at the *GNAI3* locus in the whole-genome sequencing data. (C3) Genomic PCR and direct sequencing of the PCR products. The KO allele contained a 321-bp deletion. (D1) Gene structure, sgRNA-targeted sites, and primer binding sites of *GNAO1*. (D2) IGV view of the targeted region and surrounding sequences at the *GNAO1* locus in the whole-genome sequencing data. Although the whole-genome sequencing data indicated the presence of several alleles with distinct deletions around the target site, the region around exon 6 appeared to be commonly deleted among them. (D3) Genomic PCR of the region surrounding exon 6 and sequences determined from cloned colonies. Three cloned colonies were analyzed, and all contained a 414-bp deletion, probably resulting in exon 6 skipping. The predicted amino acid sequence is shown. This alteration disrupts the G3 motif. (E1) Gene structure, sgRNA-targeted sites, and primer binding sites of GNAZ. (E2) IGV view of the targeted region and surrounding sequences at the *GNAZ* locus in the whole-genome sequencing data. (E3) Genomic PCR and sequences determined from cloned colonies. All analyzed colonies contained 355-bp deletions, and two closely related mutant junction sequences were identified. The predicted amino acid sequences deduced from the mutant alleles are C-terminally truncated by frameshift. Although the whole-genome sequencing data suggested the presence of an allele lacking this 355-bp deletion, such an allele was not detected by PCR. Thus, although the possibility that a functional *GNAZ* sequence remains cannot be excluded, this does not affect the analyses presented in this study; precise determination of the *GNAZ* genotype of this KO cell line therefore remains a subject for future investigation. (F1) Gene structure, sgRNA-targeted sites, and primer binding sites of *GNAT1*. (F2) IGV view of the targeted region and surrounding sequences at the *GNAT1* locus in the whole-genome sequencing data. (F3) Genomic PCR performed with primer pairs Fw/Rv_in and Fw/Rv, and direct sequencing of the PCR products. The KO allele contained a 1593-bp deletion. The predicted amino acid sequence deduced from the mutant allele is C-terminally truncated by frameshift. (G1) Gene structure, sgRNA-targeted sites, and primer binding sites of *GNAT2*. (G2) IGV view of the targeted region and surrounding sequences at the *GNAT2* locus in the whole-genome sequencing data. (G3) Genomic PCR performed with primer pairs Fw/Rv_in and Fw/Rv, and direct sequencing of the PCR products. The KO allele contained a 3105-bp deletion, and the predicted amino acid sequences are shown below. In addition to the 3105-bp deletion allele, the whole-genome sequencing data also suggested the presence of alleles containing an indel near exon 6 (in ENST00000679935.1) and an indel near exon 9. However, in the KO sample, no amplification product of approximately 500 bp was detected with Fw/Rv_in, whereas a faint band of >1000 bp was observed. Therefore, additional primers were designed and genomic PCR was performed. (G4) Primer Fw2 was designed further downstream of exon 9. In addition, primer Fw_in was designed to examine exon 6. Additional genomic PCR analysis of the edited *GNAT2* locus was performed using primer pairs Fw2/Rv_in and Fw_in/Rv. The Fw_in/Rv amplification product from the KO sample was approximately 200 bp smaller than that from the WT sample. Together with the WGS data visualized in IGV, this suggests that the 5′ side of exon 6 is deleted, resulting in deletion of at least amino acids 154-189.

Because this region includes the G2 motif, the gene product is predicted to be nonfunctional. Amplification of the region near exon 9 with Fw2/Rv_in was again not obtained from the KO sample, although the reason for this remains unknown. (H1) Gene structure, sgRNA-targeted sites, and primer binding sites of *GNAT3*. (H2) IGV view of the targeted region and surrounding sequences at the *GNAT3* locus in the whole-genome sequencing data. (H3) Genomic PCR was performed with the primer pairs Fw/Rv_in and Fw/Rv, and the PCR products were directly sequenced. The KO allele contained a 4736-bp deletion, and the predicted amino acid sequence is shown below. The whole-genome sequencing data also suggested the presence of an allele lacking this 4736-bp deletion. In addition, because Rv_in is located within a 240-bp deletion near one of the sgRNA target sites detected by whole-genome sequencing, no clear amplification product was obtained with Fw/Rv_in in the KO sample. Therefore, although the possibility that a functional *GNAT3* sequence remains cannot be excluded, this does not affect the analyses presented in this study; precise determination of the *GNAT3* genotype remains a subject for future investigation. (I1) Gene structure, sgRNA-targeted sites, and primer binding sites of *GNAL*. (I2) IGV view of the targeted region and surrounding sequences at the *GNAL* locus in the whole-genome sequencing data. (I3) Genomic PCR using three primer combinations, a schematic model of the mutant *GNAL* locus inferred from the PCR results, and direct sequencing of the junction region. The KO cells are considered to carry one allele with an 1867-bp deletion and another allele with an inversion of the region between the Cas9 cleavage sites. Both alterations are expected to completely delete or functionally disrupt exons 2, 3, and 4. Predicted amino acid sequences for *GNAL* isoform 1 and isoform 2 are shown below; for isoform 2, two possible mutant products are predicted, one generated by alternative splicing and the other by extension of exon 1. (J1) Gene structure, sgRNA-targeted sites, and primer binding sites of *GNAS*. (J2) IGV overview of the *GNAS* locus in the whole-genome sequencing data. (J3) Enlarged views of the sequences surrounding the two targeted regions near exon 1 and exon 13. Reads spanning the Cas9 target sites strongly suggest that an inversion has occurred between these two regions. (J4) Genomic PCR using four primer combinations and a schematic model of the mutant *GNAS* locus inferred from the PCR results, indicating an inversion between the regions containing exon 1 and exon 13. Primer pairs Fw/Rv_in and Fw_in/Rv amplified the WT cells, whereas Fw/Fw_in and Rv/Rv_in amplified the KO cells. Cloning and sequencing of colony PCR products derived from the Fw/Fw_in amplicon from the KO cells revealed a fused sequence consisting of exon 1 in the forward orientation (yellow highlight) and exon 13 in the reverse orientation (magenta highlight) joined near the

Cas9 target sites.

```
AtCnidop3a  -------------------------------MVAPHFAGFAIAASVFGSCGIVLNFTVCL
JellyOP     -------------------------------MGANITEILSGFLACVVFLSISLNMIVLI
Bovine rho  MNGTEGPNFYVPFSNKTGVVRSPFEAPQYYLAEPWQFSMLAAYMFLLIMLGFPINFLTLY

                                       89^2.56 90^2.57                113^3.28
                                   86^2.53     91^2.58 94^2.61                 117^3.32
AtCnidop3a  TYILNRRLLDASNIFILNISAGDFLYSITALPMLVTSNALGKWSFGDGGCVAYGFLTTFF
JellyOP     TFYRLRHKLAFKDALMASMAFSDVVQAIVGYPLEVFTVVDGKWTFGMELCQVAGFFITAL
Bovine rho  VTVQHKKLRTPLNYILLNLAVADLFMVFGGFTTTLYTSLHGYFVFGPTGCNLEGFFATLG

AtCnidop3a  ALGAMMNLAGAAYERYVTMCKLYESGERQFSRRKAAFLCSVLWTYALIWSIAPIFGWSSY
JellyOP     GQVSIAHLTALALDRYFTVCRPFVATAIHGSMRNAGMVIFVCWFYASFWAVLPLVGWSNY
Bovine rho  GEIALWSLVVLAIERYVVVCKPMSN--FRFGENHAIMGVAFTWVMALACAAPPLVGWSRY

                    186^ECL2
AtCnidop3a  KQEGIGTSCSTDWKS--RDVKSLSYGIVLIITCFVVPVAVILYCHIEAYKVTRKLGKQAQ
JellyOP     DVEGDGMRCSINWAD--DSPKSYSYRVCLFVFIYLIPVLLMVATYVLVQGEMKNMRGRAA
Bovine rho  IPEGMQCSCGIDYYTPHEETNNESFVIYMFVVHFIIPLIVIFFCYGQLVFTVKEAAAQQQ

AtCnidop3a  QNWGSHTRATHETLKAQKKMGEIAVVITVGFVVAWTPYTVASAIGMY---DPDLVSDVGA
JellyOP     QLFGSESEAALKNIKAEKRHTRLVFVMILSFIVAWTPYTFVAMWVSFFTKQLGPIPLYVD
Bovine rho  E--------SATTQKAEKEVTRMVIIMVIAFLICWLPYAGVAFYIFTH--QGSDFGPIFM

               293^7.40
         292^7.39   296^7.43
AtCnidop3a  SIPAYFAKSSSCYNPFIYLFMYKKLRIGMVRLLCCMKKQVYPSSSGGLTRDPTCQEIPMV
JellyOP     TLAAMLAKSSAMFNPIIYCFLHKQFRRAVLRGVCGRIVGGNAIAPSSTAVEPGQTLASGT
Bovine rho  TIPAFFAKTSAVYNPVIYIMMNKQFRNCMVTTLCCGKNPLGDDEASTTVSKTETSQVAPA

AtCnidop3a  VPVNNPL
JellyOP     AES----
Bovine rho  -------
```

**<u>Supplemental Fig. S4</u> Amino acid sequence alignment of AtCnidop3a, JellyOP, and bovine rhodopsin.**

Amino acid sequences of AtCnidop3a, JellyOP, and bovine rhodopsin (rho) are aligned using CLUSTALW (https://www.genome.jp/tools-bin/clustalw). Amino acid positions at which substitutions are introduced in this study are highlighted and their amino acid numbers are indicated. The lysine residue at position $296^{7.43}$, which is bound to the retinal chromophore, is also indicated. Amino acid residues at positions at $94^{2.61}$ and $113^{3.28}$ are colored in red.

Supplemental Table S1 the target sequences for Gs/Gq/Gi KO

| Gene | Target name | Target sequence | PAM |
|---|---|---|---|
| *GNAI1* | GNAI1_T1 | CCTTCAATGTTCCGTGACTG | TGG |
| *GNAI1* | GNAI1_T2 | AGCACTGAGTGACTACGACC | TGG |
| *GNAI2* | GNAI2_T1 | GGCCTGCTTTGGCCGCTCAA | GGG |
| *GNAI2* | GNAI2_T2 | AGCCTTGAGCGCCTATGACT | TGG |
| *GNAI3* | GNAI3_T1 | GCTGAGGCACCGTCTATTGC | TGG |
| *GNAI3* | GNAI3_T2 | CTGGAGGGCGTGCCGGGCGT | GGG |
| *GNAT1* | GNAT1_T1 | CTTCCAGCGAGTACCCGTCC | TGG |
| *GNAT1* | GNAT1_T2 | TTTGACGTTCTGCGTGTCGG | TGG |
| *GNAT2* | GNAT2_T1 | CAAGTCTTTGACGGAAAACT | TGG |
| *GNAT2* | GNAT2_T2 | ACAACTCCTATGATGATGCG | GGG |
| *GNAT2* | GNAT2_T3 | AGAAGAGGCCGCAGTCCTTG | AGG |
| *GNAT2* | GNAT2_T4 | ACGTCTCTTAGCCTCGTCTG | TGG |
| *GNAT3* | GNAT3_T1 | GGCAACCTTACGCACCTAAG | TGG |
| *GNAT3* | GNAT3_T2 | AATCCACCTAAACCGCAGTT | TGG |
| *GNAO1* | GNAO1_T1 | CTCGGTGGGCTGGTAGTCGG | CGG |
| *GNAO1* | GNAO1_T2 | CCGTGACGTCCTCGAAGCAA | TGG |
| *GNAZ* | GNAZ_T1 | GTTGTGGAAGTCGATCCTGA | GGG |
| *GNAZ* | GNAZ_T2 | AGAGGTCAGAGCGCAAAAAG | TGG |
| *GNAL* | GNAL_T1 | CAGCAAGACGACGGAAGACC | AGG |
| *GNAL* | GNAL_T2 | CAGGGGAATTGCTTGATCGA | GGG |
| *GNAS* | GNAS_T1 | CCTCGGTCTTACTGTTCCCG | AGG |
| *GNAS* | GNAS_T2 | AGAGCAGCTCGTACTGACGA | AGG |

Supplemental Table S2 The primer sequences used for genotyping

| Gene | Primer name | Sequence | Amplicon size (in WT) | |
|---|---|---|---|---|
| *GNAI1* | GNAI1_Fw | TTCTTTACATCAAAGCCTTGC | 692 bp | |
| *GNAI1* | GNAI1_Rv | TGGCAACACCTTCAGCTCTTCATGT | | |
| *GNAI2* | GNAI2_Fw | GGTTTTAGGGCAAGTCTCATCC | 506 bp | |
| *GNAI2* | GNAI2_Rv_in | ATAAGCCACCTGCCTGTCATTC | | |
| *GNAI2* | GNAI2_Rv | CCTGTGTACTCAGGGAAGCAGA | | |
| *GNAI3* | GNAI3_Fw | TCGGATATCCGGTTCTTCTGG | 485 bp | |
| *GNAI3* | GNAI3_Rv | CCGGAGTCACACCCTCTACAG | | |
| *GNAO1* | GNAO1_Fw | AATTTTTGTGCCCCCTCGGGAGTAG | 729 bp | |
| *GNAO1* | GNAO1_Rv | AGCACCTGGTCATAGCCGCTG | | |
| *GNAZ* | GNAZ_Fw | ACGCCGCGAAATCAAGCTGC | 601 bp | |
| *GNAZ* | GNAZ_Rv | GCCGCTGAGCTCCACACAGA | | |
| *GNAT1* | GNAT1_Fw | CAACTCGGGAGGTCCCCGTC | 332 bp | |
| *GNAT1* | GNAT1_Rv_in | TGAGTGTGGTCATGGCGCGT | | |
| *GNAT1* | GNAT1_Rv | AGGCGCTGGGATACTGTGGG | | |
| *GNAT2* | GNAT2_Fw | TGGCCTGTGCTCTTTAAGCTGC | 647 bp | |
| *GNAT2* | GNAT2_Rv_in | AGCAGGCATTAGATGCAGGAGAGC | | 813 bp |
| *GNAT2* | GNAT2_Fw2 | GCCTTCCCTTCAGTTATGTTCGTGC | | |
| *GNAT2* | GNAT2_Rv | GCAGTCAGGGCAAACCCAAGG | 537 bp | |
| *GNAT2* | GNAT2_Fw_in | AAGTGGGAGAGGAAAAGGGGGAGAA | | |
| *GNAT3* | GNAT3_Fw | AGCAAAACGTTGGATGGTTGTC | 371 bp | |
| *GNAT3* | GNAT3_Rv_in | AGCTAGGTGAGTTGCAGGGAAG | | |
| *GNAT3* | GNAT3_Rv | TTTGTAAATAGTTGGATATTCCATTGGT | | |
| *GNAL* | GNAL_Fw | AAAACACCGAAGAGACCAGACCATC | 625 bp | |
| *GNAL* | GNAL_Rv | CCGATTATCAGCCTTCCCCATACAC | | |
| *GNAL* | GNAL_Fw_in | TGGCCAACCCTGAAAACCAATTTCG | | |
| *GNAS* | GNAS_Fw | CTGGCTCTCTGTGTCTCTCGCTCTT | 930 bp | |
| *GNAS* | GNAS_Rv_in | CCTTCTGCAGCTGCTTCTCGATCTT | | |
| *GNAS* | GNAS_Fw_in | TACTGCTACCCTCATTTCACCTGCG | 261 bp | |
| *GNAS* | GNAS_Rv | AAAGGGGGTTTCGCAAAATCACTCG | | |

**<u>Supplemental Data</u> Amino acid sequences of Nluc inserted human GsαL and Venus fused human Gγ1 for BRET-based Gs Protein Dissociation Assay.**

Nluc inserted human GsαL:

MGCLGNSKTEDQRNEEKAQREANKKIEKQLQKDKQVYRATHRLLLLGAGESGKSTIVKQMRILHVNGFNGEGGEE

DPQAARSNSDGEKATKVQDIKNNLKEAIETIVAAMSNLVPPVELANPENQFRVDYILSVMNSGGGGTRSVFTLEDFV

GDWRQTAGYNLDQVLEQGGVSSLFQNLGVSVTPIQRIVLSGENGLKIDIHVIIPYEGLSGDQMGQIEKIFKVVYPVDD

HHFKVILHYGTLVIDGVTPNMIDYFGRPYEGIAVFDGKKITVTGTLWNGNKIIDERLINPDGSLLFRVTINGVTGWRL

CERILATGGGGSVPDFDFPPEFYEHAKALWEDEGVRACYERSNEYQLIDCAQYFLDKIDVIKQADYVPSDQDLLRCR

VLTSGIFETKFQVDKVNFHMFDVGGQRDERRKWIQCFNDVTAIIFVVASSSYNMVIREDNQTNRLQEALNLFKSIWN

NRWLRTISVILFLNKQDLLAEKVLAGKSKIEDYFPEFARYTTPEDATPEPGEDPRVTRAKYFIRDEFLRISTASGDGRH

YCYPHFTCAVDTENIRRVFNDCRDIIQRMHLRQYELL

Black: GsαL, Red: Nluc, Blue: linker

Venus fused human Gγ1:

MVSKGEELFTGVVPILVELDGDVNGHKFSVSGEGEGDATYGKLTLKLICTTGKLPVPWPTLVTTLGYGLQCFARYPD

HMKQHDFFKSAMPEGYVQERTIFFKDDGNYKTRAEVKFEGDTLVNRIELKGIDFKEDGNILGHKLEYNYNSHNVYI

TADKQKNGIKANFKIRHNIEDGGVQLADHYQQNTPIGDGPVLLPDNHYLSYQSALSKDPNEKRDHMVLLEFVTAAG

ITLGMDELYKGSAGTMPVINIEDLTEKDKLKMEVDQLKKEVTLERMLVSKCCEEFRDYVEERSGEDPLVKGIPEDKN

PFKELKGGCVIS

Black: Gγ1, Red: Venus, Blue: linker
